# *Drosophila* Neuroblast Selection Gated by Notch, Snail, SoxB and EMT Gene Interplay

**DOI:** 10.1101/783241

**Authors:** Badrul Arefin, Farjana Parvin, Shahrzad Bahrampour, Caroline Bivik Stadler, Stefan Thor

## Abstract

In the developing *Drosophila* central nervous system neural progenitor (neuroblast; NB) selection is gated by lateral inhibition, controlled by Notch signalling and proneural genes. However, proneural mutants only display partial NB reduction, indicating the existence of additional genes with proneural activity. In addition, recent studies reveal involvement of key epithelial-mesenchymal transition (EMT) genes in NB selection, but the regulatory interplay between Notch signalling and the EMT machinery is unclear. We find that the SoxB gene *SoxNeuro* and the Snail gene *worniou* are integrated with the Notch pathway, and constitute the missing proneural genes. Notch signalling, the proneural, *SoxNeuro*, and *worniou* genes regulate key EMT genes to orchestrate the NB specification process. Hence, we uncover an expanded lateral inhibition network for NB specification, and demonstrate its link to key players in the EMT machinery. Because of the evolutionary conservation of the genes involved, the Notch-SoxB-Snail-EMT network may control neural progenitor selection in many other systems.

## INTRODUCTION

The embryonic *Drosophila melanogaster* (*Drosophila*) central nervous system (CNS) has been a central model system for addressing the genetic mechanisms controlling neural progenitor specification. The *Drosophila* CNS can be separated into the brain and the ventral nerve cord, which are generated from the head neurogenic and ventral neurogenic regions, respectively (Figure 1A). The process of neural progenitor (denoted neuroblasts in *Drosophila*; NBs) generation has been most extensively studied in the ventral neurogenic regions, where some ~60 bilateral NBs and a smaller number of midline NBs form in each segment of the neuroectoderm, during early to mid-embryogenesis (Figure 1A) (Birkholz et al., 2013; Bossing et al., 1996; Schmid et al., 1999; Schmidt et al., 1997; Urbach et al., 2016; Wheeler et al., 2009).

**Figure 1.**
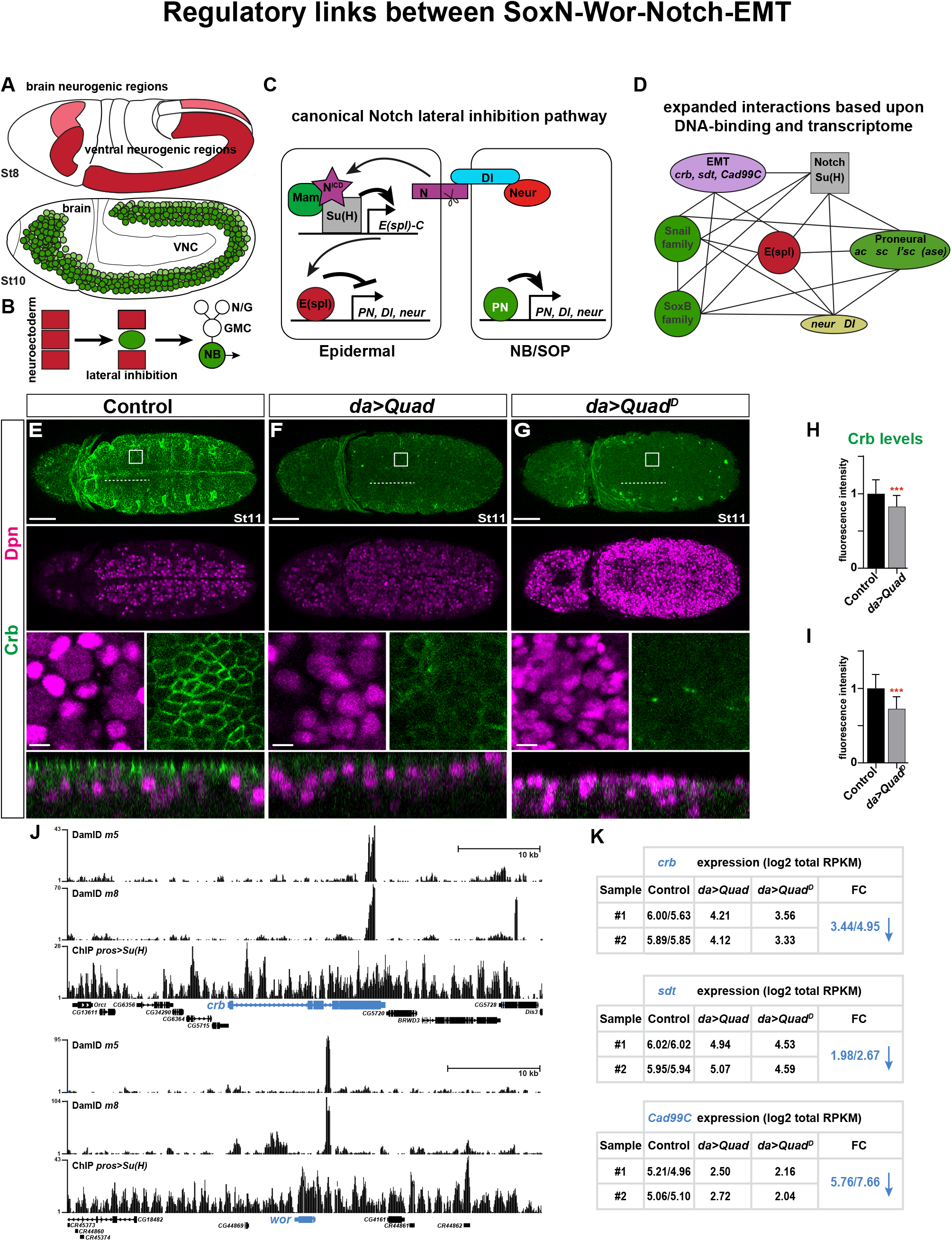
Regulatory links between SoxN-Wor-Notch-EMT. (A) Cartoon of *Drosophila* embryos, showing the brain and the ventral neurogenic regions (upper panel, St8) from where the brain and the ventral nerve cord (VNC) originate, respectively (bottom, St10). (B) Cartoon of NB selection (green) in the neuroectoderm, by the process of lateral inhibition, followed by NB delamination, and lineage progression (GMC=ganglion mother cell; N/G=neuron/glia). (C) Canonical Notch lateral inhibition pathway (PN=proneural). (D) Genetic interactions, as well as novel interactions based upon DNA-binding and RNA-seq analysis (see main text, and Supplemental Tables 1 and 2, for details). (E-G) Whole mount embryos showing expression of Dpn (NBs) and Crb, at St11; dorsal view, anterior to the left. Combinatorial co-misexpression of four early NB factors triggers extensive ectopic NB generation and reduced Crb expression. Control = *da-Gal4>+. da>UAS-Quad* = *UAS-ase, UAS-wor, UAS-SoxN, UAS-Kr. da>UAS-Quad^D^* = *UAS-ase-Vp16, UAS-wor-EnR, UAS-SoxN-EnR, UAS-Kr*. Boxes and dotted lines on the whole embryo represent magnified panels and orthogonal view orientations below, respectively. Scale bars: whole embryo = 50μm, magnified panels = 10μm. (H-I) Quantification of Crb levels within the boxed regions (fluorescence intensity, mean in control was set to one; Student’s two-tailed T-test; ***p≤0.001; n≥60 boxes, n≥4 embryos; mean+/-SD). (J) DamID-seq analysis of E(spl)m5 and E(spl)m8, from *UAS-E(spl)m5* and *UAS-E(spl)m8* St9-16 embryos, as well as ChIP-seq analysis of Su(H) from *pros-Gal4/UAS-Su(H)*, St9-16, embryos [based upon data from (Bivik et al., 2016)]. DNA-binding analysis reveals binding of all three proteins to the *crb* and *wor* genes. (K) RNA-seq analysis from St9-16 embryos. *crb, sdt* and *Cad99C* are downregulated by ≥2-fold in both *da-Gal4>UAS-Quad* and *da-Gal4>UAS-Quad^D^*.

NBs are selected in the neuroectoderm by a process denoted lateral inhibition, after which they delaminate and rapidly commence generating neural lineages (Figure 1B). The Notch pathway plays a central role during the lateral inhibition process (Bray, 2016; Bray and Gomez-Lamarca, 2018; Hori et al., 2013; Kopan and Ilagan, 2009; Perdigoto and Bardin, 2013; Schweisguth, 2015). In the canonical Notch lateral inhibition pathway, the Delta (Dl) ligand binds to the Notch receptor, and ubiquitination of Dl, by the E3 ligase Neuralized (Neur) (Deblandre et al., 2001; Lai et al., 2001; Pavlopoulos et al., 2001; Yeh et al., 2001), results in Dl endocytosis, which promotes the trans-activation of Notch. This triggers Notch cleavage, releasing the Notch intracellular domain (NICD), which enters the nucleus and forms a tripartite complex with the DNA binding factor Suppressor of Hairless [Su(H)] and the co-factor Mastermind (Mam) (Hori et al., 2013; Kopan and Ilagan, 2009). This complex activates the *Enhancer-of-split Complex* (*E(spl)-C*) of transcription factors (TFs); founding members of the HES gene family of bHLH transcriptional repressors (Bailey and Posakony, 1995; Delidakis and Artavanis-Tsakonas, 1992; Knust et al., 1992; Lecourtois and Schweisguth, 1995). The TFs encoded by the *E(spl)-C* in turn repress the bHLH proneural genes within the *achaete-scute complex* (*AS-C*) i.e., *achaete* (ac), *scute* (sc) and *lethal of scute* (*l’sc*) (Giagtzoglou et al., 2003). In addition, the *E(spl)-C* also represses the *neur* and *Dl* genes, as well as the *asense* (*ase*) gene; a proneural-related bHLH gene located within the *AS-C* (Heitzler et al., 1996; Miller and Posakony, 2018; Miller et al., 2014). Downregulation of the proneural genes inhibits NB formation, instead promoting epidermal differentiation (Jimenez and Campos-Ortega, 1990). Cells with low Notch activation instead continue proneural expression, which in turn promotes proneural, *neur* and *Dl* expression, promoting NB fate, while continuing to present Dl to neighbouring cells, hence promoting their epidermal fate (Figure 1C). The lateral inhibition process underlying the selection of peripheral sensory organ precursors (SOPs) has also been the subject of intense studies and in many key aspects mirrors the NB selection process (Schweisguth, 2015). However, in spite of the substantial body of work that have helped decode the Notch pathway and the lateral inhibition process, there are several key outstanding issues.

First, while it is clear that the proneural genes are critical for NB generation, previous studies demonstrated that even deletion of the entire proneural gene complex (*AS-C*) only results in the failure to generate a subset of NBs (Jimenez and Campos-Ortega, 1990). This has prompted the speculation of the existence of additional proneural genes (Skeath and Carroll, 1994; Skeath and Thor, 2003). But what is the identity of these missing proneural genes? Although not typically considered as proneural genes, members of the SoxB family to some extent qualify for this role. The central role of the SoxB family in NB selection has been, in part, obscured by the genetic redundancy between the two SoxB genes *SoxNeuro* (*SoxN*) and *Dichaete* (D). However, *SoxN/D* double mutants do show severe reduction of NB numbers (Buescher et al., 2002; Overton et al., 2002). Another gene family with some proneural-like activity is the Snail family; *snail*, (*sna*), *escargot* (*esg*) and *worniou* (*wor*), which act redundantly during CNS development (Ashraf et al., 1999; Ashraf and Ip, 2001; Cai et al., 2001). While, *sna, esg, wor* triple mutants do not show an apparent reduction in early NB numbers (Ashraf et al., 1999), *wor* mutants, also heterozygous for *sna* and *esg*, display a loss of NBs at later stages (Bahrampour et al., 2017). In addition, co-misexpression of a set of NB factors, which included Wor and SoxN, was shown to be sufficient for generating ectopic NBs broadly in the developing embryonic ectoderm, and even in developing wing discs (Bahrampour et al., 2017). This raises the issue of how *SoxN/wor* co-misexpression can completely override the negative, anti-NB, input from Notch signalling, and whether or not they may be considered to constitute the missing proneural genes.

Second, NB delamination is akin to an epithelial-mesenchymal transition (EMT) process (An et al., 2017; Doe, 1992; Hartenstein and Campos-Ortega, 1984; Kraut and Campos-Ortega, 1996; Simoes et al., 2017). Moreover, studies reveal that the EMT and apical polarity gene *crumbs* (*crb*) modulates Notch signalling (Das and Knust, 2018; Richardson and Pichaud, 2010). Intriguingly, recent findings show that Neur not only regulates Dl endocytosis, but also regulates Crb endocytosis, by controlling Stardust (Sdt) stability (another EMT and apical polarity gene; Pals1 in mammals) (Perez-Mockus et al., 2017). These findings suggest an intimate interplay between Notch signalling and some key EMT genes. However, the mechanistic intersection between the Notch pathway and the EMT genes is unclear.

To address these outstanding issues, we analysed the effects of loss- (LOF) and gain-of-function (GOF) of the Notch pathway, the proneural genes, the EMT gene *crb*, the SoxB family gene *SoxN* and the Snail family gene wor, upon NB generation and upon the expression of the same four entities, in the developing *Drosophila* embryo. We find that *SoxN* and *wor* are both necessary and sufficient for NB selection. Strikingly, *SoxN/wor* coexpression can generate massive numbers of NBs even in an *AS-C* mutant background. These GOF and LOF effects leads us to propose that *SoxN* and *wor* constitute the missing proneural genes. We also find that the Notch pathway, the proneural, *SoxN, wor* and EMT genes, are involved in elaborate cross-regulation. These findings greatly expand the lateral inhibition cascade and also link it mechanistically to the EMT network. Our results further support the notion that NB delamination is an EMT-like process, albeit with several modifications.

Recent studies in mammals of the neuroepithelial-to-Radial Glia Cell transition suggest that this process also bears resemblance to an EMT-like process, and that most, if not all of the genes acting in *Drosophila* may be involved in the related mammalian process (Camargo Ortega et al., 2019; Itoh et al., 2013; Singh et al., 2016; Zander et al., 2014). Hence, neural progenitor selection may be an evolutionary conserved modified EMT-like process, involving a conserved cassette of Notch-Snail-SoxB-EMT-gene interplay.

## RESULTS

### Transcriptome analysis reveals that NB factors regulate EMT and asymmetry genes

We recently found that combinatorial misexpression of a number of NB TFs could trigger ectopic NB generation, both in the embryonic ectoderm and the developing wing discs (Bahrampour et al., 2017). These TFs included SoxN, Wor and Ase, which were particularly potent in combination also with Kruppel (Kr) i.e., *UAS-ase, -SoxN, -wor, -Kr* (denoted *UAS-Quad*). In addition, a combination of dominant versions of three of these TFs i.e., *UAS-ase-Vp16, -SoxN-EnR, -wor-EnR, -Kr* (denoted *UAS-Quad-dominant*; *Quad^D^*) was also highly potent (Bahrampour et al., 2017). To further address the effects of these TFs, we co-misexpressed the *Quad* and *Quad^D^*, using the *da-Gal4* driver line, which expresses both maternally and ubiquitously zygotically (Wodarz et al., 1995). We found that both the *da>Quad* and *da>Quad^D^* misexpression triggered massive ectopic NB generation, evident at St11 by the expression of Dpn (Figure 1E-G). Orthogonal views revealed ectopic NBs not only in the underlying cell layers, but also in the upper neuroectodermal layer (Figure 1E-G; bottom). These extremely potent effects of early NB TFs may have gone unnoticed before due to posttranscriptional control of these transgenes. However, these recently generated *UAS* transgenes were based upon synthetic and codon-optimized cDNAs (Bahrampour et al., 2017). These constructs furthermore omitted the endogenous 5’ and 3’ UTR, to thereby reduce the possible unwanted negative influence of miRNA control. They also contained optimized start-ATG regions, and were site-specifically integrated in the genome, using phiC31 landing site technology (Bischof et al., 2007). Together with the combinatorial approach, these improvements apparently uncover a dramatic potency of these TFs in generating NBs. Intriguingly, co-misexpression of these early NB TFs can apparently completely override the Notch lateral inhibition pathway.

To gain further insight into the possible connection between SoxN, Wor and Ase with Notch signalling, we analysed previous RNA-seq data (Bahrampour et al., 2017), with RNA prepared from stage (St) 8-16 embryos, for *da-Gal4* driving the two quadruple *UAS* combinations. We previously found that both *Quad* and *Quad^D^* triggered extensive ectopic expression of both neuronal and glia markers, *elav* and *repo*, respectively, indicating that the ectopic NBs generated underwent full neuronal differentiation. In addition, a number of “driver” cell cycle genes i.e., *Cyclin E, E2f1* and *stg* are upregulated (Bahrampour et al., 2017). Focusing on the Notch pathway, and we also observed upregulation of *l’sc* and *neur* (Table S1).

In addition to these, mostly logical, effects on gene expression, we also found that both the *da>Quad* and *da>Quad^D^* triggered more than two-fold change (≥2FC) down-regulation of a number of genes involved in the EMT process, including *crb, sdt*, and *Cad99C* (Figure 1K and Table S1). The repression of *crb* was confirmed by staining for Crb, which revealed down-regulation of Crb in *da>Quad* and *da>Quad^D^* (Figure 1E-I). In contrast to the effects on the EMT genes *crb, sdt*, and *Cad99C*, genes in the Par-complex (Par-C) i.e., *baz/par-3, par-6* and *aPKC*, as well as genes in the Scribble-complex (Scrib-C) i.e., *scrib, dlg1* and *l(2)gl* (*lgl*) were not altered by ≥2FC (Table S1). Moreover, genes involved in asymmetric division of NBs i.e., *miranda, inscuteable, prospero* and *partner of numb* (*pon*) were in fact ≥2FC upregulated (Table S1) (Bahrampour et al., 2017).

### DNA-binding analysis indicates extensive interplay between the Notch pathway, *SoxN, wor*, and EMT genes

An extensive number of gene-specific DNA-binding studies have unravelled clear direct transcriptional regulatory links between components within the Notch signalling pathway, showing that Su(H), Sc, L’Sc and Ac bind directly to the *ac, E(spl)-complex, neur* and/or *Su(H)* genes (Table S2) (Bailey and Posakony, 1995; Barolo et al., 2000; Cave et al., 2005; Kageyama et al., 1997; Krejci and Bray, 2007; Lecourtois and Schweisguth, 1995; Liu and Posakony, 2014; Miller and Posakony, 2018; Nellesen et al., 1999; Oellers et al., 1994; Tietze et al., 1992; Van Doren et al., 1994). Moreover, genome-wide DNA-binding approaches involving ChIP-seq of Su(H), as well as DamID-seq analysis of DamID-fusions to E(spl)m5, E(spl)m8 and Ase, have confirmed these links (Table S2) (Bernard et al., 2010; Bivik et al., 2016; Djiane et al., 2013; Housden et al., 2013; Krejci et al., 2009; Southall and Brand, 2009).

We re-analysed our recent genome-wide ChIP-seq and DamID-seq data for Su(H), E(spl)m5 and m8 binding in the embryo (Bivik et al., 2016), and observed binding to all of the genes in the Snail and SoxB families i.e., *wor, sna, esg, D* and *SoxN* (Figure 1J and Table S2). Moreover, previous DamID-seq of Ase identified direct binding to wor, *sna* and *D* (Southall and Brand, 2009), and studies have revealed binding of Sna and D to many Notch pathway components (Table S2) (Aleksic et al., 2013; Negre et al., 2011; Southall and Brand, 2009). Finally, the Notch pathway TFs Su(H), m5, m8, Ase, as well as Sna and D, also bind to several key EMT genes, including *crb, sdt* and *Cadherin 99C* (*Cad99C*) (Figure 1J and Table S2).

In summary, the RNA-seq and DNA-binding data indicate extensive regulatory interplay between the Notch pathway, proneural, SoxN and Wor genes, as well as the EMT and asymmetry genes, beyond the canonical lateral inhibition pathway (Figure 1D).

### SoxN and Wor expression is controlled by the Notch pathway

To determine the extent to which the aforementioned possible regulatory interplay is indeed functionally involved in NB selection, we analysed Notch pathway, *SoxN, wor*, and *crb* mutants and misexpression, scoring for NB numbers and protein/reporter expression levels.

First, we focused on the Notch pathway. As anticipated, *Notch* and *neur* mutants displayed a striking increase in the number of NBs generated, as revealed by Dpn and Ase staining (Figures 2A-B, 2P, S1A-B, S1P). Conversely, simultaneous removal of all three proneural genes (*ac, sc* and *l’sc*, as well as the related *ase* gene), by deletion of the *achaete-scute complex* (*AS-C*) genomic region, resulted in a reduction of NB numbers (Figure S1C, S1P). Activation of the Notch pathway, by pan-embryonic expression of the Notch intracellular domain [*da>NICD;* (Go et al., 1998)] or a stabilized version of *E(spl)m8* [*da>m8^CK2^*; (Bivik et al., 2016)] both resulted in reduction of NBs (Figure 2C-E, 2P). Misexpression of proneural genes (*da>l’sc*) did not however alter NB numbers (Figure S1D-E, S1P).

**Figure 2.**
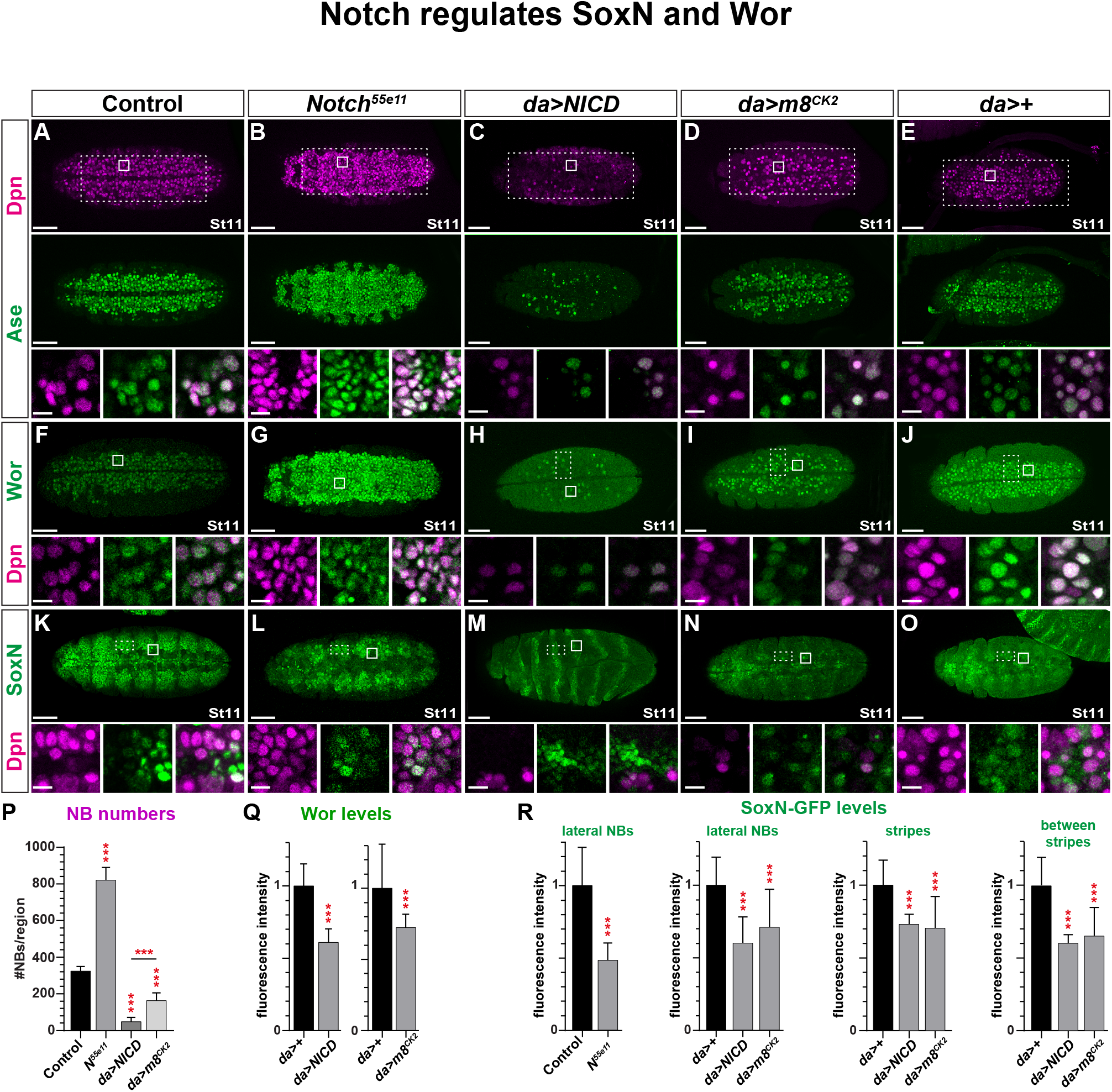
Notch regulates SoxN and Wor. (A-O) Expression of Dpn, Ase, Wor and *SoxN-GFP* in control, Notch mutant and Notch pathway activated embryos, at St11; ventral view, anterior to the left. (A-B) In *Notch^55e11^* mutants, excessive numbers of NBs are generated. (C-E) In contrast, *da-Gal4/UAS-NICD*, and *da-Gal4/UAS-m8^CK2^* embryos show reduction of NB numbers. (A-J) Ase and Wor expression shows similar expression profile, and co-localizes with Dpn in NBs, in both mutant and misexpression embryos (magnified panels). (K-O) In control, *SoxN-GFP* has a broader expression pattern than Dpn, Ase and Wor, being expressed both in NBs and in neighboring cells. *SoxN-GFP* expression responds in a complex manner to both *Notch^55e11^* mutant and *da-Gal4>UAS-NICD* and *da-Gal4/UAS-m8^CK2^* misexpression. Embryos in (K-L) are homozygous for *SoxN-GFP*, while (M-O) are heterozygous. (P) Quantification of NB numbers inside the large dashed rectangles in (A-E) (Student’s two-tailed T-test; ***p<0.001; n>4 embryos; mean+/-SD). (Q-R) Quantification of Wor levels in NBs, and *SoxN-GFP* levels in lateral NBs, as well as in stripes and between stripes (fluorescence intensity; Student’s twotailed T-test; ***p≤0.001; n≥30 boxes, n≥3 embryos; mean+/-SD). Scale bars: whole embryo = 50μm, magnified panels = 10μm.

Next, we turned to Notch pathway regulation of Wor and SoxN expression. In the wild type, Wor expression mirrored that of Dpn and Ase, being expressed in most, if not all, NBs as they emerge during St8-10 (Figure 2F, S1F). Mutating or activating the Notch pathway resulted in changes in Wor expression that followed Dpn and Ase (Figure 2F-J, 2Q, S1F-J, S1Q). Although proneural misexpression (*da>l’sc*) did not trigger extra NBs it did affect Wor expression within NBs. However, surprisingly, Wor expression was slightly reduced (Figure S1I-J, S1Q).

*SoxN* expression is broader and earlier than that of Dpn, Ase and Wor, and commences in the presumptive neurogenic regions already at syncytial blastoderm stage (Cremazy et al., 2000). Subsequently, *SoxN* is expressed broadly in neuroectodermal cells, and becomes elevated in NBs as they form (Cremazy et al., 2000). To monitor *SoxN* expression we utilized a *SoxN-GFP* transgenic fosmid line, where GFP is fused to the C-terminus of SoxN within the context of the *SoxN* genomic region (Sarov et al., 2016). Surprisingly, given the supernumerary NBs formed, we found that *Notch* and *neur* mutants displayed reduction of *SoxN-GFP* expression in the neuroectoderm and in NBs (Figure 2K-L, 2R, S1K-L, S1R). More logically, *da>NICD* and *da>m8^CK2^* expression, which greatly reduces NB numbers, also resulted in reduced *SoxN-GFP* expression (Figure 2M-O, 2R). *da>NICD* and *da>m8^CK2^* triggered a more pronounced *SoxN-GFP* stripe expression in the ectoderm, although expression in both in the stripes and in-between-stripes was reduced (Figure 2M-O, 2R). In proneural mutants (*AS-C*), *SoxN-GFP* was down-regulated in NBs, while proneural misexpression (*da>l’sc*) resulted in *SoxN-GFP* being upregulated (Figure S1M-O, S1R).

In summary, the well-established roles of the Notch pathway and the proneural genes results in logical effects upon Wor expression in NBs, which largely mirror Dpn and Ase expression. The expression of *SoxN-GFP* displays a more complex picture, and both Notch pathway GOF and LOF results in reduced expression. Logically however, proneural LOF and GOF reveal that they are positive regulators of *SoxN-GFP* expression.

### *SoxN* and *wor* regulate the Notch pathway

Next, we addressed the possible reciprocal connection between *SoxN, wor* and the Notch pathway. To this end we focused on *E(spl)* expression as a readout of Notch pathway activation, because they are well-established Notch targets (Bailey and Posakony, 1995; Cooper et al., 2000; Krejci and Bray, 2007; Lecourtois and Schweisguth, 1995; Nellesen et al., 1999; Wurmbach et al., 1999), and reporter transgenes using *E(spl)* enhancers-promoters have been demonstrated to faithfully report upon Notch signalling (Castro et al., 2005; Kramatschek and Campos-Ortega, 1994; Lai et al., 2000; Nolo et al., 2000). We previously used an *m8-GFP* reporter to detect Notch activity (Ulvklo et al., 2012), and using this reporter, we scored Notch activity in the developing neuroectoderm in *SoxN* and *wor* mutants, as well as in single and double misexpression embryos. We also scored the number of NBs generated, as detected by Dpn and Ase staining.

The Snail family shows genetic redundancy (Ashraf et al., 1999; Ashraf and Ip, 2001; Cai et al., 2001), but even *sna,esg,wor* triple mutants did not show any apparent reduction of early NB numbers (Ashraf et al., 1999). In contrast, our previous studies showed that *wor* mutants did display reduced NB numbers at later stages (Bahrampour et al., 2017). Perhaps a reason for this single-gene effect was that the mutant background was sensitized for *sna* and *esg*, by placing the *wor^4^* allele, a strong *wor* allele (Ashraf et al., 2004), over a genomic deletion that removes wor, *sna* and *esg* (*Df(2L)ED1054*; referred to as *wor^Df^* herein). Hence, this mutant combination is homozygous for *wor* and heterozygous for *sna* and *esg*, but we refer herein to this allelic combination as “wor mutants”. Similarly, within the SoxB family, *SoxN* is expressed more broadly, in lateral NBs, while *D* is restricted to medial NBs (Buescher et al., 2002; Cremazy et al., 2000; Ma et al., 2000; Nambu and Nambu, 1996; Overton et al., 2002; Soriano and Russell, 1998). In line with the broader expression of *SoxN*, previous studies showed that *SoxN* was the strongest gene regarding NB generation (Overton et al., 2002), and we previously observed reduced NB numbers in *SoxN* mutants (Bahrampour et al., 2017). We therefore focused on *SoxN* within the SoxB family, and used a nonsense mutation allele (*SoxN^NC14^*) (Chao et al., 2007) over the genomic deletion *Df(2L)Exel7040* (referred to as *SoxN^Df^* herein).

As anticipated from previous studies, we observed a reduction in NB numbers in both *SoxN* and *wor* mutants (Figure 3A-C, 3F). The *UAS-Quad* and -*Quad^D^* combinatorial expression, which includes *SoxN* or *wor*, can trigger massive ectopic NB generation in the embryonic ectoderm or wing imaginal discs (Figure 1E-G) (Bahrampour et al., 2017; Bahrampour et al., 2019). However, we did not previously test the effects of single *SoxN* or *wor* misexpression from an early *Gal4* driver. Strikingly, in both *SoxN* and *wor* single misexpression, driven by *da-Gal4*, we observed generation of extra NBs (Figure 3F). These effects were stronger for the dominant-negative transgenic versions, *SoxN-EnR* and *wor-EnR* (denoted *SoxN^D^* and *wor^D^* herein) (Figure 3F). Co-misexpression of *SoxN* and wor, either as wild type or dominant versions (*da>Double* and *da>Double^D^*), resulted in even more extensive NB generation (Figure 3D-F). These extremely potent effects of *SoxN/wor* misexpression may have gone unnoticed before, due to posttranscriptional control of these transgenes. However, our recently generated synthetic codon-optimized cDNAs (Bahrampour et al., 2017) apparently uncovers a dramatic potency of *SoxN/wor* in generating NBs.

**Figure 3.**
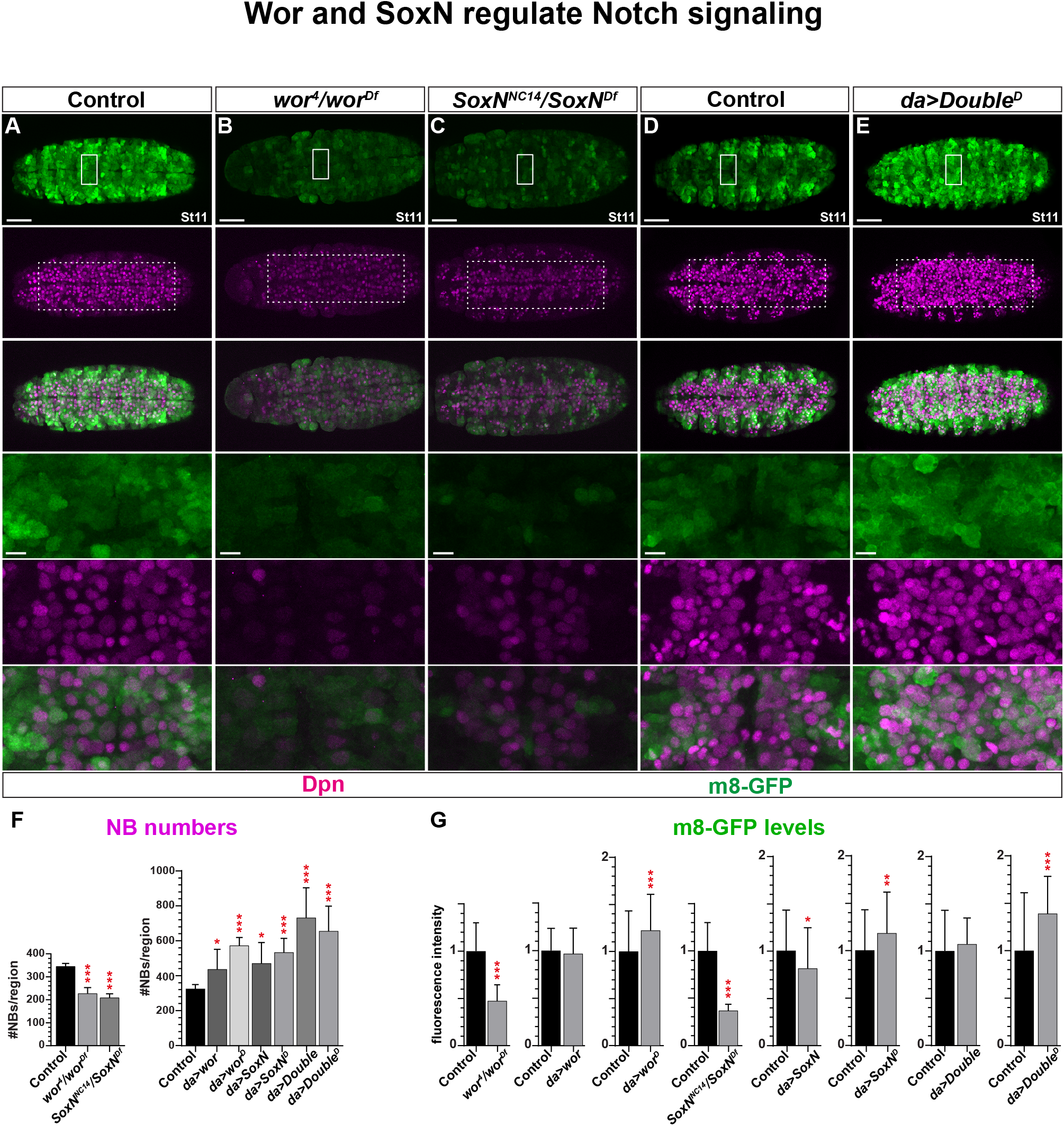
*SoxN* and *wor* regulate Notch signalling. (A-E) NB numbers (Dpn+) are reduced in *wor^4^/wor^Df^* and *SoxN^NC14^/SoxN^Df^* mutants, whereas they are increased in *da-Gal4>Double^D^* co-misexpression embryos. *m8-GFP* reporter expression is reduced in *wor^4^/wor^Df^* and *SoxN^NC14^/Sox*.mutants, whereas it is upregulated in *da-Gal4>Double^D^*. Embryos in (A-C) are homozygous for *m8-GFP*, while (D-E) are heterozygous. (F) Quantification of NB numbers (NBs quantified inside large dashed rectangle outline in (A-E); Student’s two-tailed T-test; ***p≤0.001; **p≤0.01; *p≤0.05; n≥30 sections, n>3 embryos; mean+/-SD). Single misexpression of wor, *SoxN, wor^D^* and *SoxN^D^*, as well as co-misexpression (*Double* and *Double^D^*), all result in a significant increase in NB numbers. (G) Quantification of m8-GFP levels (fluorescence intensity; Student’s two-tailed T-test; ***p≤0.001; **p≤0.01; *p≤0.05; n≥30 sections, n≥4 embryos; mean+/-SD). Scale bars: whole embryo = 50μm, magnified panels = 10μm. Stage 11.

Turning to *m8-GFP* expression, surprisingly, in spite of the reduction in NB numbers in *SoxN* and *wor* mutants, a typical Notch^ON^ phenotype, we observed reduction of *m8-GFP* expression in both cases (Figure 3A-C, 3G). Conversely, in spite of supernumerary generation of NBs in all misexpression embryos, a typical Notch^OFF^ phenotype, we observed a mixed effect on *m8-GFP* reporter expression, with *wor^D^, SoxN^D^* and *Double^D^* showing elevated *m8-GFP* expression, while *SoxN* displayed reduced expression (Figure 3D-E, 3G). Hence, we find an intriguing dichotomy between the effects of *wor* and *SoxN* on NB generation versus *m8-GFP* expression.

### *SoxN* and *wor* can generate supernumerary NBs even in proneural mutants

While proneural genes are important for NB selection, previous studies (Jimenez and Campos-Ortega, 1990) and our results herein (Figure S1C, S1P), demonstrate that even removal of all proneural gene activity (deletion of the entire *AS-C* complex) does not result in complete absence of NBs. This has prompted the speculation of the existence of additional proneural genes (Skeath and Carroll, 1992; Skeath and Thor, 2003). Similar to the proneural genes, *wor* and *SoxN* mutants also display reduction of NB numbers (Figure 3F). This prompted us to address the possible redundancy between these genes. Strikingly, we find that *AS-C;wor* double mutants display a near complete loss of NBs, significantly below the numbers observed in each single mutant (Figure 4A-B, 4F).

**Figure 4.**
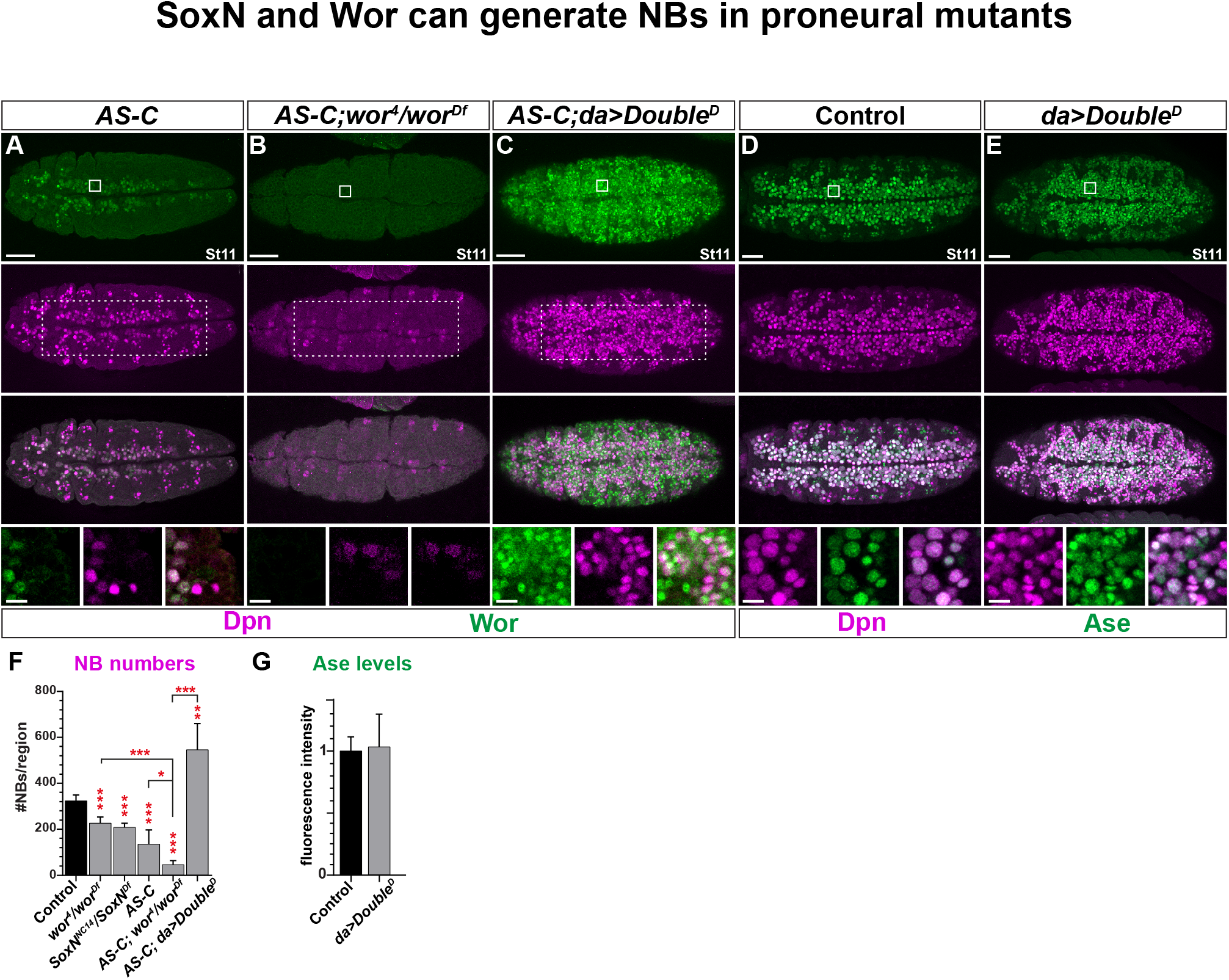
*SoxN* and *wor* rescue proneural mutants. (A-C) NB numbers are reduced in proneural mutants (AS-C), and is even more reduced in *AS-C;wor^4^/wor^Df^*. However, *da-Gal4* driven misexpression of *UAS-Double^D^* in the *AS-C* mutant background still generates supernumerary NBs. (D-E) Co-misexpression of *SoxN^D^* and *wor^D^* (*da-Gal4>UAS-Double^D^*) generates excessive numbers of Dpn/Ase positive NBs (magnified panels-bottom). (F) Quantification of NB numbers inside the large dashed rectangles in (A-C) (Student’s two-tailed T-test; ***p≤0.001; **p≤0.01; n≥3 embryos; mean+/-SD) (note, data for control, *wor* and *SoxN* mutant numbers were reproduced from Figure 3F). (G) Quantification of Ase levels (fluorescence intensity; Student’s two-tailed T-test; n≥30 sections, n≥3 embryos; mean+/-SD). Scale bars: whole embryo = 50μm, magnified panels = 10μm. Stage 11.

*SoxN* and *wor* misexpression, and in particular co-misexpression, resulted in the generation of extensive numbers of ectopic NBs (Figure 3F). We analysed how this affected Ase, and indeed observed ectopic expression of Ase, which accompanied Dpn, albeit without increasing Ase levels in Dpn cells (Figure 4D-E, 4G).

These two findings prompted us to test if *SoxN/wor* co-misexpression could generate supernumerary NBs even in the absence of proneural gene activity. To this end we leaned on the strong effect of the *SoxN^D^* and *wor^D^* (*da>Double^D^*), and co-misexpressed them in the *AS-C* deletion background. Strikingly, we observed that while *AS-C* displayed a reduction of NB numbers, co-misexpression of *SoxN/wor* (*da>Double^D^*) in *AS-C* mutants still resulted in the generation of massive numbers of ectopic NBs, as evident by Dpn expression (Figure 4A, 4C, 4F).

Thus, not only do *SoxN* and *wor* fit the definition as bona fide proneural genes by their LOF and GOF effects, showing that they are both necessary and sufficient for NB generation, they can also generate NBs in a genetic background completely lacking *AS-C* proneural gene activity.

### *crb* and the Notch pathway regulate each other

Previous studies revealed that *crb* is important for Notch signalling in the developing eye disc (Richardson and Pichaud, 2010). In the embryonic ectoderm, studies suggest that Crb acts to stabilize the Notch receptor in the membrane, thereby enhancing Notch signalling, evident by increased NB numbers in *crb* mutants (Das and Knust, 2018).

We analysed Dpn in *crb* mutants, and as anticipated observed increased NB numbers (Figure 5A-B, 5K). Conversely, *crb* overexpression resulted in fewer NBs (Figure 5C-D, 5L). The effect of *crb* on Notch pathway activation was not previously directly tested. To address this we again used *m8-GFP* as readout. We observed down-regulation of *m8-GFP* in *crb* mutants (Figure 5A-B, 5I). Surprisingly, *crb* overexpression also resulted in *m8-GFP* down-regulation (Figure 5C-D, 5J). These results point to that a balance in Crb levels is important for proper Notch signalling.

**Figure 5.**
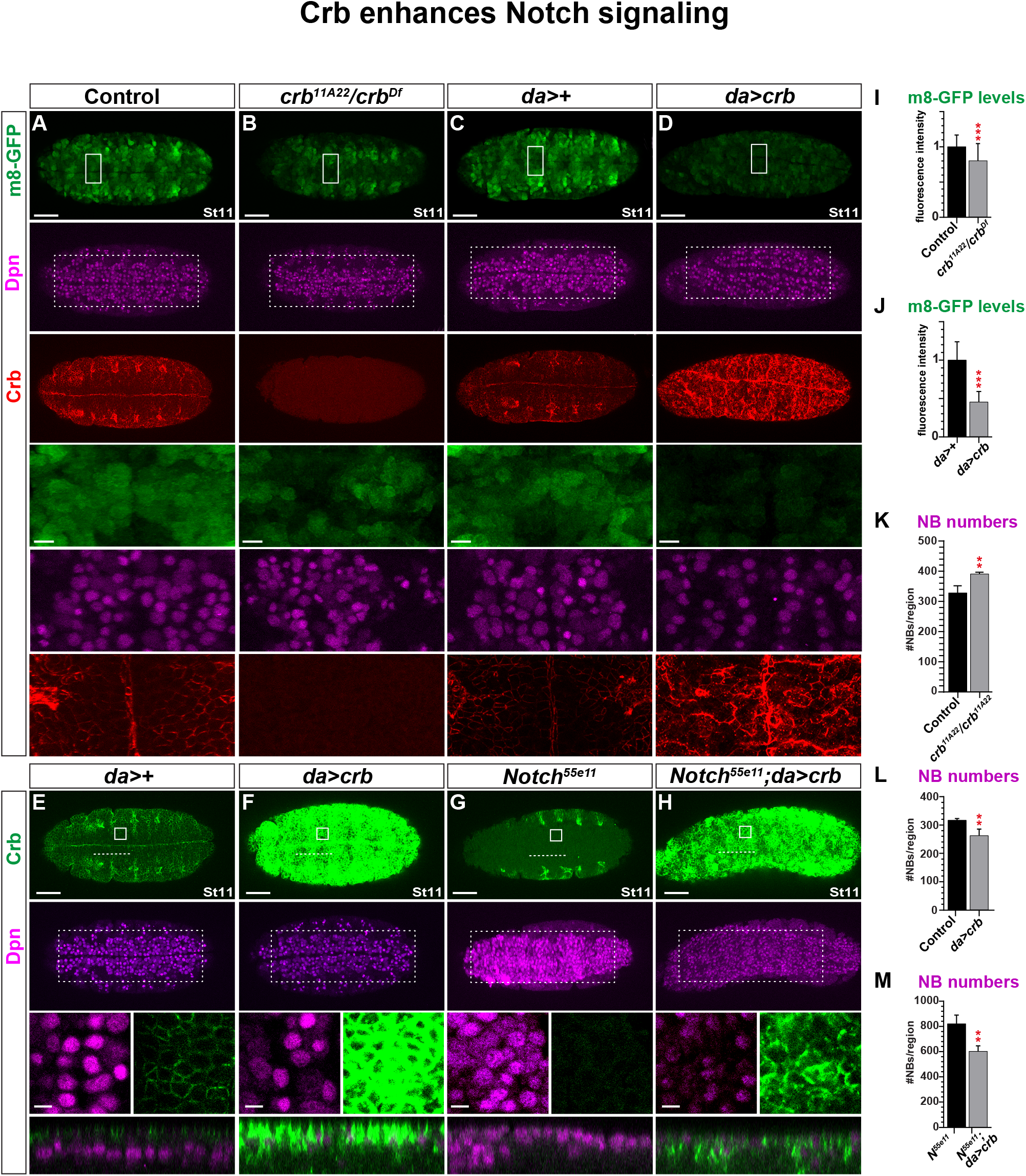
*crb* enhances Notch signalling. (A-D) NB numbers are increased *crb* mutants (*crb^11A22^/crb^Df^*) and reduced in *crb* overexpression (*da-Gal4>UAS-crb*). However, *m8-GFP* reporter expression is reduced in both *crb* mutant and *crb* overexpression embryos. (E-F) *crb* overexpression (*da-Gal4>UAS-crb*) results in reduced NB numbers and elevated Crb expression. (G-H) Notch mutants (*Notch^55e11^*) display increased NB numbers, but *crb* overexpression (*da-Gal4>UAS-crb*) in *Notch* mutants reduces NB numbers. Boxes depicted on whole embryos are magnified below; dotted lines are shown as orthogonal views below. In control, NBs (Dpn+ cells) are primarily observed delaminated from the overlying neurectodermal cell layer. In contrast, in *Notch* mutants, NBs are formed already in the overlying neurectodermal cell layer. Cross-rescue of *Notch* with *da>crb* reduces the number of NBs forming, but NBs are still observed in the overlying ectodermal layer. Embryos in (A-D) are heterozygous for *m8-GFP*. (I-J) Quantification of m8-GFP levels (fluorescence intensity; Student’s two-tailed T-test; ***p≤0.001; **p≤0.01; n≥30 regions; n≥3 embryos; mean+/-SD). (K-M) Quantification of NB numbers inside the large dashed rectangles in (A-D, G-H) (Student’s two-tailed T-test; **p≤0.01; n≥3 embryos; mean+/-SD). Scale bars: whole embryo = 50μm, magnified panels = 10μm. Stage 11.

The possible regulation of Crb by Notch signalling was not hitherto addressed. We found striking downregulation of Crb in both *Notch* and *neur* mutants (Figure S2A-C, S2G-H). Conversely, we observed strong upregulation of Crb in both *da>NICD* and *da>m8^CK2^* (Figure S2D-F, S2I-J).

These results prompted us to test if *crb* transgenic expression could at least in part rescue *Notch* mutants. The *Notch^55e11^* allele used in our study is a *copia-like* transposon element insertion in the *Notch* gene (Kelley et al., 1987), and has been described as either amorph or hypomorph (Campos-Ortega, 1983; Tian et al., 2004). Considering that it may be a hypomorph, we attempted to rescue *Notch^55e11^* mutants by transgenic expression of *crb*. Strikingly, we observed partial rescue of *Notch* mutants by transgenic expression of *crb*, evident by a reduction of NB numbers in *Notch^55e11^;da>crb* embryos, when compared to *Notch^55e11^* alone (Figure 5E-H, 5M).

Next, we addressed Crb regulation by the proneural genes. Somewhat surprisingly, we observed that *AS-C* mutants, which display a reduction of NB numbers, showed reduced Crb expression (Figure S3A-B, S2E). We also observed that misexpression of proneural genes (*da>l’sc*) reduced Crb expression (Figure S3C-D, S3F).

These findings reveal that Crb and Notch pathway cross-regulation is critical for proper Notch signalling and NB generation.

### *SoxN, wor* and *crb* regulate each other

Finally, we turned to the interplay between wor, *SoxN* and *crb*. Surprisingly, in spite of the reduction in NB numbers in *wor* and *SoxN* mutants, a typical Notch^ON^ phenotype, we observed reduced Crb expression in both mutants (Figure 6A-C, 6G). Less surprising were the *wor* and *SoxN* misexpression effects, which generate extra NBs, revealing reduced Crb expression for *da>wor^D^, -SoxN* and -*SoxN^D^*, as well as both of the double *UAS* combinations (Figure 6D-G).

**Figure 6.**
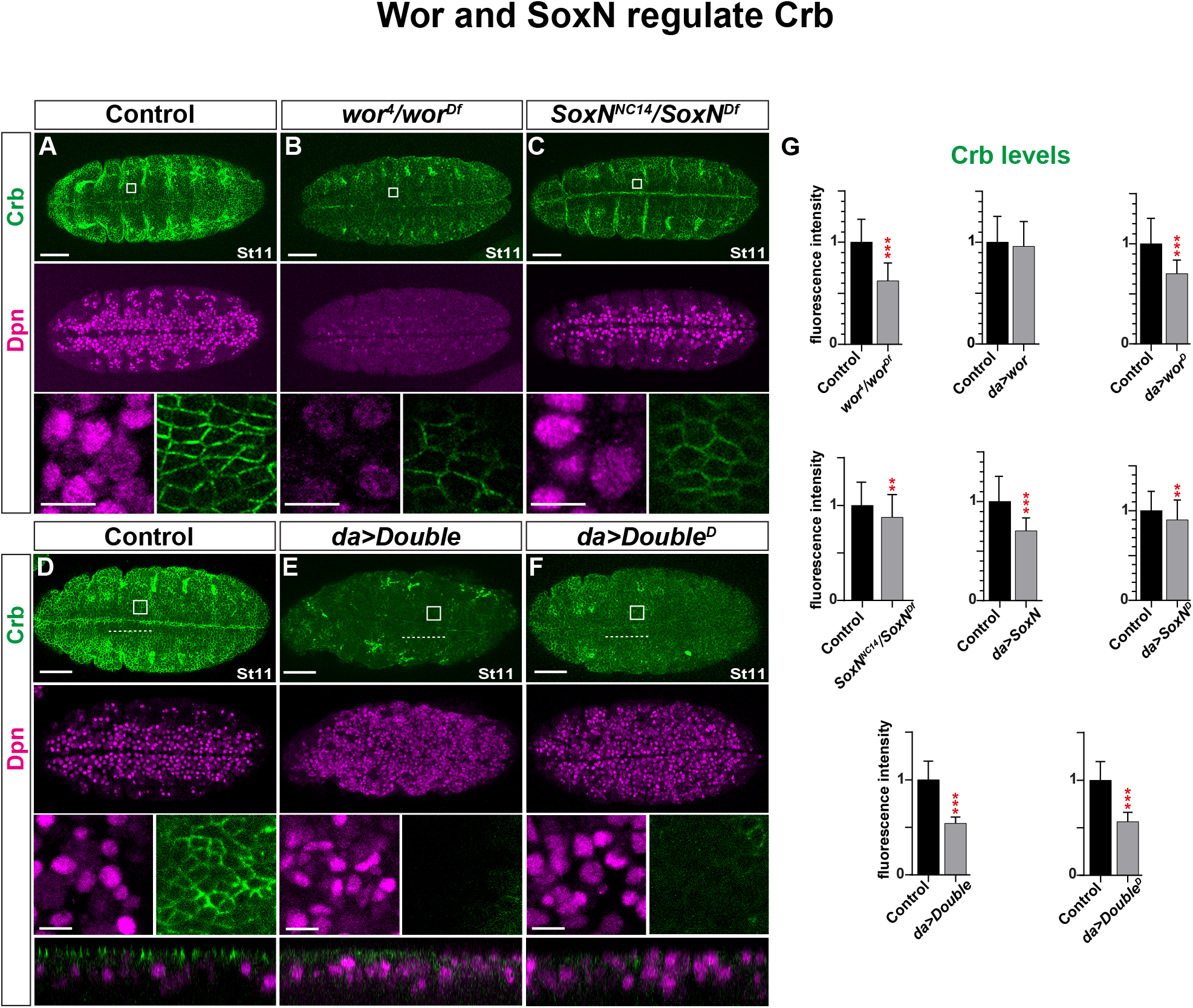
*SoxN* and *wor* regulate Crb expression. (A-F) Crb expression is reduced in *wor^4^/wor^Df^* and *SoxN^NC14^/SoxN^Df^* mutants and co-misexpression (*da-Gal4/UAS-Double* and *UAS-Double^D^*). Boxes depicted on whole embryos are magnified below; dotted lines are shown as orthogonal views below. (G) Quantification of Crb levels (Student’s two-tailed T-test; ***p≤0.001; **p≤0.01; n≥30 sections, n≥3 embryos; mean+/-SD). Single misexpression of *SoxN, wor^D^* and *SoxN^D^*, as well as co-misexpression (*Double* and *Double^D^*), all result in significant reduction of Crb expression. Scale bars: whole embryo = 50μm, magnified panels = 10μm. Stage 11.

We also performed the reciprocal experiments, and found that *crb* mutants show a reduction of Wor expression in NBs, as well as of *SoxN-GFP* expression in the neuroectoderm and in NBs (Figure S4A-H, S4Q-R). Overexpression of *crb* also reduced *SoxN-GFP* expression, both in NBs and in the neuroectoderm, while Wor expression in NBs was unaffected (Figure S4I-P, S4Q-R).

## DISCUSSION

### *SoxN* and *wor*: the missing proneural genes?

Based upon previous studies, and our findings herein, *SoxN* is necessary for NB generation (Buescher et al., 2002; Overton et al., 2002). Regarding the Snail family, previous studies did not find apparent reduction of NBs numbers in *sna, esg, wor* triple mutants (Ashraf et al., 1999). However, these studies focused on early stages of neurogenesis, and may not have covered the complete span of NB formation. Analysing embryos at St11, after most, if not all, NBs have formed, we find that *wor* mutants i.e., *wor^4^* placed over a deletion that simultaneously removes one gene copy of wor, *sna* and *esg*, do indeed display a significant reduction in NB numbers [herein; (Bahrampour et al., 2017)]. In addition, while *AS-C* and *wor* mutants both show a partial loss of NBs, we find that *AS-C;wor* double mutants display a significantly more severe loss than either single mutant alone. Moreover, misexpression of *SoxN* or wor, from our novel optimized transgenes, reveals a dramatic sufficiency for these genes in generating ectopic NBs in the ectoderm [herein; (Bahrampour et al., 2017)]. Strikingly, *SoxN/wor* co-misexpression can generate extensive numbers of ectopic NBs even in a genetic background lacking any *AS-C* proneural gene activity. Finally, both *SoxN* and *wor* are regulated by, and regulate, the Notch pathway (see below). Based upon these findings we propose that *SoxN* and *wor* constitute the missing proneural genes.

What are the connections between the Notch pathway, *SoxN* and wor? Notch signalling, based upon *Notch* and *neur* mutants, as well as *NICD* and *m8* misexpression, represses *SoxN* and Wor expression inside NBs. However, the reciprocal connection i.e., between *SoxN/wor* and the Notch pathway, as measured by *m8-GFP* expression, is complex and in some instances dichotomous. Specifically, *SoxN* and *wor* mutants display fewer NBs; a Notch^ON^ effect, but reduced *m8-GFP* expression; a Notch^OFF^ effect. Similarly, misexpression of *SoxN^D^* and *wor^D^* trigger more NBs; a Notch^OFF^ effect, but increased *m8-GFP* expression; a Notch^ON^ effect.

Previous studies found that the proneural genes i.e., *ac, sc* and *l’sc*, are upstream of *wor* (Ashraf and Ip, 2001). In line with this, in *AS-C* mutants we observe loss of Wor cells (NBs) and reduction of Wor expression levels in the NBs still generated. Similarly, we previously found that *ase* misexpression increased Wor expression in NBs (Bahrampour et al., 2017). Previous studies revealed that *SoxN* mutants show reduced Ase expression in NBs, and that misexpression of *SoxN* could activate Wor expression, whereas neither *wor* or *ase* GOF or LOF affected SoxN expression in NBs (Bahrampour et al., 2017). In addition, *SoxN* mutants were found to display loss of *ac, l’sc* and Wor expression (Overton et al., 2002). In summary, *SoxN*, which is expressed in the entire early neuroectoderm, acts upstream of the proneural genes, while proneural genes act upstream of wor. However, this SoxN->proneural->wor regulatory flow is complex e.g., while both *SoxN* and *wor* misexpression can trigger NB generation, *l’sc* does not have this potency. Moreover, while *SoxN* expression may occur first, both *AS-C* and *wor* appear to be important for maintained and elevated *SoxN* expression in NBs. Hence, *SoxN, AS-C* and *wor* appear to be involved in a mutually reinforcing interplay, which ensures robust NB selection once the Notch pathway balanced is tipped (Figure 7A).

**Figure 7.**
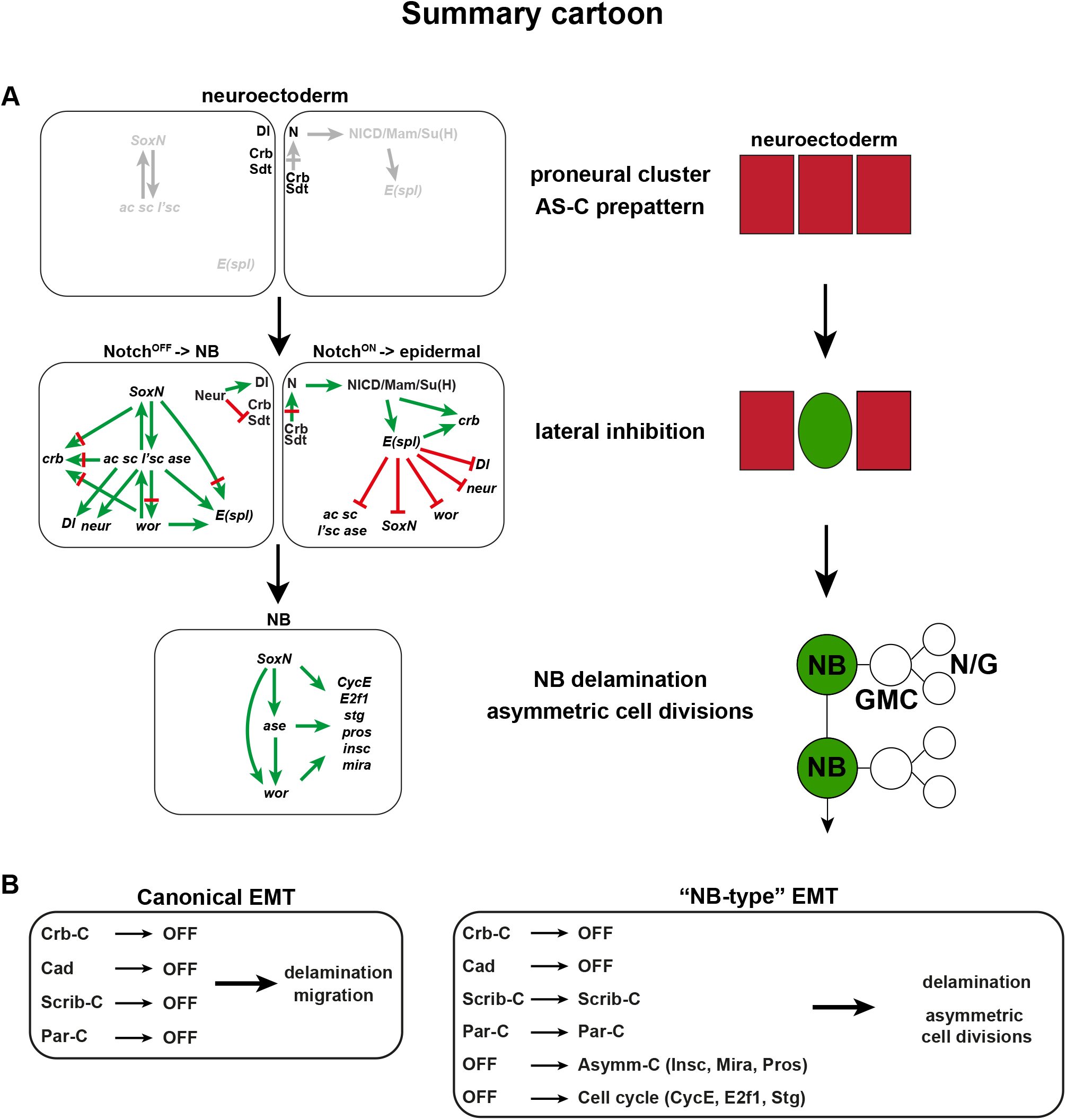
Summary cartoon. (A) An expanded lateral inhibition cascade acts during *Drosophila* NB selection and delamination. In this model, the canonical Notch pathway is also regulated by the SoxB gene *SoxN* and the Snail gene wor, as well as by the modulation of Notch receptor activation by the Crb/Sdt complex. (Top) In the early neuroectoderm, there is simultaneous and weak expression of SoxN, AS-C and E(spl) genes, and weak activation of Notch signalling. (Middle) Subsequently, the lateral inhibition process triggers elevated expression of *SoxN* and *AS-C* in the Notch^OFF^ cells, while the Notch^ON^ cells elevate *E(spl)* expression. The outcome of these events is activation of *wor* and *ase* in the Notch^OFF^ cells, and downregulation of *crb*, as well as other EMT genes, leading to the delamination of the NB. (Bottom) The combined action of SoxN, Ase and Wor ensures activation of the cell cycle and asymmetric genes, resulting in repetitive rounds of asymmetric cell division. (B) In the canonical EMT process, the Crb, Scrib and Par complexes, as well as Cadherins, are downregulated, leading to delamination from the epithelial sheet and cell migration. In the “NB-type” EMT, only the Crb complex and Cadherins are down-regulated, while the Scrib and Par complexes remain expressed. This triggers delamination from the epithelial sheet, similar to canonical EMT, but retains apico-basal cell polarity in the NB. In addition, the outcome of the expanded lateral inhibition process (A) results in the expression of proneural genes, SoxN, Wor and Ase, which triggers the expression of asymmetric genes (e.g. *insc, mira, pros*) and cell cycle driver genes (e.g. *CycE, E2f1, stg*(*cdc25*)), the outcome of which is to drive repetitive rounds of asymmetric cell divisions of the NB. Hence, “NB-type” EMT represents a modification of the canonical EMT process, ensuring that delamination is followed by the unique behaviour of NBs during CNS lineage progression.

DNA-binding studies for the factors studied herein, and analysis of related family members (D, Sna, Ase), suggest that the elaborate transcriptional interplay between all of the aforementioned transcription factors/co-factors i.e., NICD/Su(H)/Mam, E(spl), proneural, SoxN, Wor, and their respective genes may result from direct transcriptional regulation (Figure 7A, Table S2).

### Interplay of *crb*, Notch signalling, *SoxN* and *wor* during NB selection

During mammalian EMT, the Crb complex, which contains the Crb, Sdt (mammalian Pals1) and Patj proteins, plays a key role (Dongre and Weinberg, 2019; Lamouille et al., 2014). In *Drosophila, crb* and *sdt* also control epithelial polarity in a number of tissues (Bachmann et al., 2001; Bulgakova and Knust, 2009; Campbell et al., 2009; Grawe et al., 1996; Hong et al., 2001; St Johnston and Ahringer, 2010; Tepass, 1996; Tepass et al., 1990; Thompson et al., 2013), while *Patj* only acts in ovaries (Penalva and Mirouse, 2012).

Recent studies revealed that Crb stabilizes Notch, and accordingly *crb* mutants show more NBs [herein; (Das and Knust, 2018)]. We furthermore find that *crb* overexpression results in fewer NBs, and, strikingly, that *crb* overexpression can partly rescue Notch mutants. The reduction of NB numbers in *crb* mutants is logically accompanied by reduced *m8-GFP* expression, while, surprisingly, *crb* misexpression also triggered reduced *m8-GFP* expression. We find that Notch signalling activates Crb expression, evident by downregulation of Crb in *Notch* and *neur* mutants, and upregulation of Crb in *NICD* and *m8* misexpression. Hence, with the exception of *crb* misexpression on *m8-GFP*, a clear-cut interplay between canonical Notch signalling and *crb*/Crb emerges (Figure 7A). However, this interplay would constitute a runaway loop, with Notch activating *crb*, and Crb supporting Notch activation. Perhaps our finding that *crb* overexpression reduces *m8-GFP* points to a nebulous brake pedal in this loop.

Regarding the connection between *crb* with *SoxN, wor*, and the proneural genes, we also find clear interplay, albeit with a reverse logic for mutants versus misexpression. Specifically, misexpression of *SoxN, wor* or proneural genes, which generates more NBs, with encompassing delamination, also results in reduced Crb expression. However, surprisingly, *SoxN, wor* and proneural mutants, which display fewer NBs, also show reduced Crb expression, underscoring the balancing act of these gene regulations (Figure 7A).

In addition, further complexity regarding the role of *crb* stems from recent findings revealing that Neur, an E3 ligase critical for Dl endocytosis (Deblandre et al., 2001; Lai et al., 2001; Pavlopoulos et al., 2001; Yeh et al., 2001), also controls the stability of Sdt, and thereby affects Crb protein levels (Perez-Mockus et al., 2017). It is tempting to speculate that Notch^OFF^ cells (NBs), which maintain proneural gene expression and hence activate *neur* expression, will have increased Neur, and hence increased endocytosis of Sdt, and thereby decreased Crb levels, leading to reduced Notch receptor activation. Because Neur also increases endocytosis of Dl, high Neur levels would help drive Notch activation in the neighbouring cells (epidermal cells). Moreover, since Notch (NICD and m8^CK2^) activates Crb expression and represses *neur* expression, this should ensure more Crb in the Notch^ON^ cells (epidermal cells), thereby further supporting Notch activation. By these mechanisms, the transcriptional regulation of *crb/neur* gene expression and the stability/endocytosis control of Crb/Sdt/Dl levels and localization, and thereby Notch activation, acts as a hitherto undiscovered loop providing additional thrust to the lateral inhibition decision (Figure 7A).

Similar to the TF interplay described above, the gene-specific and/or genome-wide DNA-binding studies indicate that the gene regulation of *crb* and *neur* may be mediated by direct transcriptional regulation of the Notch pathway TFs (NICD/Su(H)/Mam, E(spl), proneural), as well as the SoxB and Snail family TFs (Table S2).

### NB selection and delamination; an EMT-like process

EMT has been extensively studied in mammals, and has revealed roles for the Crb, Scribble (Scrib) and Par complexes, as well as for Notch signalling and the Snail family (Dongre and Weinberg, 2019; Lamouille et al., 2014). More recently, the SoxB family has also been implicated in EMT (Acloque et al., 2011), in particular with respect to malignant variants of EMT occurring in various cancer forms (Mladinich et al., 2016; Xu and Yang, 2017).

Previous studies, and our findings herein, demonstrate that the majority of these genes and protein complexes also play key roles also during *Drosophila* NB selection and delamination. This supports the notion that NB selection and delamination could be viewed as an EMT-like process. However, the NB-type EMT differs from canonical EMT in several aspects. In canonical EMT, all apical polarity complexes (Crb, Par and Scrib complexes) and Cadherins are turned off, and there is no activation of asymmetry genes. Hence, delamination is followed by symmetric cell division and cell migration. In contrast, in “NB-type EMT”, while, similarly, the Crb complex and Cadherins are turned off, the Par complex (*baz/par-3, par-6, aPKC*) and the Scribble complex (*scrib, dlg, lgl*) remain expressed, and asymmetric genes e.g., *mira, insc* and *pros* are turned on. In addition, within NBs, *SoxN, wor* and *ase* activate key cell cycle driver genes i.e., *Cyclin E* and *stg*, and repress expression of the cell cycle inhibitor *dacapo*, while conversely, the Notch pathway represses *Cyclin E* and activates *dacapo* (Ashraf and Ip, 2001; Bahrampour et al., 2017; Bivik et al., 2016). These gene expression changes results in NB delamination, but retains apico-basal polarity in the NB, and ensures repetitive rounds of asymmetric cell divisions, generating the unique features of CNS lineages (Figure 7B).

Based upon our findings and those previously published, a model emerges wherein SoxB acts early to govern neuroectodermal competence, intersecting with the early transient wave of proneural gene expression in the proneural clusters. SoxN and proneural genes engage in interplay with the Notch-mediated lateral inhibition process, which is gated also by Crb-Sdt-Neur membrane-localized control of Notch receptor activity and Delta ligand endocytosis. The outcome of these interactions is that early NBs become Notch^OFF^, elevate their SoxN and proneural expression, as well as activate Wor and Ase expression. This results in the downregulation of a subset of EMT genes (i.e., Crb complex and Cadherins), while the Scrib and Par complexes are maintained. The combined action of SoxN, proneural, Wor and Ase triggers activation of asymmetric cell division genes and cell cycle driver genes, the outcome of which is NB delamination, followed by asymmetric cell divisions and lineage generation. In contrast, the surrounding Notch^ON^ cells continue expressing *E(spl)* genes, downregulate the *SoxN*, proneural, *neur* and *Dl* genes. This results in the continued expression of the Crb, Scrib and Par complexes, as well as failure to activate wor, *ase*, asymmetric and cell cycle genes. The combined effect of these regulatory decisions is that these cells remain in the ectoderm and do not divide (Figure 7A).

### Neural progenitor selection and delamination; a conserved EMT-like process?

In mammals, the neuroepithelial-to-Radial Glia Cell transition (NE-RGC) is in many aspects analogous to the NB selection and delamination process in *Drosophila*. Intriguingly, recent studies suggest that NE-RGC can perhaps also be viewed as an EMT-like process, although in this case the process has been modified even further, and the RGC retains an apical connection throughout neurogenesis, and undergoes interkinetic nuclear migrations (Camargo Ortega et al., 2019; Itoh et al., 2013; Singh et al., 2016; Zander et al., 2014). Strikingly, in two recent studies the NE-RGC transition was found to involve the mouse Snail and Scratch factors, both of which are members of the Snail family (Itoh et al., 2013; Zander et al., 2014). Other players in the NB-selection program outlined above also play key roles in the early development of the mammalian CNS and in the NE-RGC transition (Cau and Blader, 2009; Dongre and Weinberg, 2019; Lamouille et al., 2014; Masek and Andersson, 2017; Reiprich and Wegner, 2015; Sarkar and Hochedlinger, 2013), although the direct comparison of gene function and cell behaviour becomes nebulous. Hence, it would appear that the neuroectoderm->neural progenitor selection and delamination process has undergone several evolutionary modifications, perhaps becoming less and less akin to a canonical EMT process in more derived animals, such as mammals. Nevertheless, it is tempting to speculate that several of the basic principles of EMT are utilized in the mammalian NE-RGC process, and that viewing it as such may be helpful for future studies.

## MATERIALS AND METHODS

### Fly stocks

#### Mutants

*wor^4^* (Bloomington Drosophila Stock Center, stock number BL#25170). *wor^Df^* = *Df(2L)ED1054* (BL#24112). *SoxN^NC14^* (BL#9938). *SoxN^Df^* = *Df(2L)Exel7040* (BL#7811). *neur^1^* (BL#4222). *neur^11^* (BL#2747). *Notch^55e11^* (BL#28813). *AS-C* = *Df(1)BSC530* (BL#25058). *crb^11A22^* (BL#3448). *crb^Df^* = *Df(3R)BSC638* (BL#25728).

#### *UAS* transgenes

*UAS-ase-myc (28E), UAS-Kr-V5* (*53B*); *UAS-wor-FLAG* (*65B*), *UAS-wor-FLAG-EnR* (*65B*), *UAS-SoxN-V5* (*89E*), *UAS-SoxN-V5-EnR* (*89E*) (Bahrampour et al., 2017). *UAS-l’sc* = *UAS-l(1)sc* (BL#51670). *UAS-m5^CK2^* and *UAS-m8^CK2^* (Bivik et al., 2016). *UAS-NICD* (Go et al., 1998), (provided by Spyros Artavanis-Tsakonas, Harvard Medical School, Boston, MA, USA). *UAS-crb* (BL#5444).

#### *Gal4* drivers

*pros-Gal4* on chromosome III (provided by Chris Q. Doe, University of Oregon, OR, USA). *da-Gal4* on chromosome II and III (BL#55849).

#### Reporter lines

*E(spl)HLH-m8-GFP* (Castro et al., 2005) (provided by James Posakony, UCSD, La Jolla, CA, USA). *SoxN-GFP* (Vienna Drosophila Resource Center #318062).

Mutants or lethal lines were maintained over GFP- or YFP-marked balancer chromosomes. *OregonR* or *Gal4/+* were used as controls. Embryos were staged according to Campos-Ortega and Hartenstein (Campos-Ortega and Hartenstein, 1985). All *Drosophila* experiments were performed at +25°C.

### Immunohistochemistry

Immunohistochemistry was performed as previously described (Bahrampour et al., 2017). Primary antibodies were: Guinea pig anti-Dpn (1:100), rat anti-Dpn (1:500) (Ulvklo et al., 2012). Chicken anti-GFP (1:1,000; cat.no. ab13970, Abcam, Cambridge, UK). Rabbit anti-Ase (1:1,000; provided by Yuh-Nung Jan, UCSF, San Francisco, CA, USA). Rat anti-Wor (1:1,000; cat.no. ab196362, Abcam, Cambridge, UK). Mouse mAb anti-Crb (Cq4; 1:50) (Developmental Studies Hybridoma Bank, Iowa City, IA, USA).

### RNA-sequencing analysis

The *da-Gal4/UAS-Quad* and *da-Gal4/Quad^D^* RNA-seq data was previously generated (Bahrampour et al., 2017) (GEO accession number GSE103660).

### DamID-seq and ChIP-seq analysis

Peak analysis of previously generated DamID-seq and ChIP-seq data (Bivik et al., 2016) (GEO accession number GSE137195), for *pros-Gal4-driven* and non-driven *UAS-m5^CK2^* and *UAS-m8^CK2^*, as well as *pros-Gal4/UAS-Su(H)*, was conducted using DNASTAR Seqman NGN software (v.12.2) (Madison, WI, USA) for sequence assembly. Normalization was performed with RPKM. Qseq was used for peak detection and the wig-files were aligned to genome assembly dm6 on the UCSC genome browser for visualization (Karolchik et al., 2014). DNA-binding data for other TFs, gene-specific or genome-wide, were mined based on previous studies (Aleksic et al., 2013; Bailey and Posakony, 1995; Barolo et al., 2000; Bernard et al., 2010; Bivik et al., 2016; Cave et al., 2005; Djiane et al., 2013; Housden et al., 2013; Kageyama et al., 1997; Krejci et al., 2009; Krejci and Bray, 2007; Lecourtois and Schweisguth, 1995; Liu and Posakony, 2014; Miller and Posakony, 2018; Negre et al., 2011; Nellesen et al., 1999; Oellers et al., 1994; Southall and Brand, 2009; Tietze et al., 1992; Van Doren et al., 1994).

### Confocal imaging, fluorescence intensity measurements and image preparation

Zeiss LSM700 and LSM800 confocal microscopes were used for imaging and confocal stacks were visualized using LSM software. Identical confocal setting was used to scan the control and experimental embryos. Dpn+ cells were used for NB number quantification. ImageJ was used for both NB number quantification and protein intensity measurement (Schindelin et al., 2012). Mean pixel intensity X area of the individual cell was referred to as fluorescence intensity. Adobe Photoshop and Adobe Illustrator CC 2018 were used for compiling of the Images and graphs.

### Statistical analysis

GraphPad Prism 8.0.1 (244) was used for the statistical analysis. Two-tailed Student’s T-test was performed unless stated otherwise (for specific statistical test used, see text and figures). Significance of p<0.05 is stated with one star (*), p<0.01 with two stars (**), p<0.001 with three stars (***). Microsoft Excel for Office 365 MSO (16.0.11601.20174) was used for data compilation.

## AUTHOR CONTRIBUTIONS

BA, FP, SB and ST designed the experiments. BA and FP performed all experiments. BA, FP, SB and CBS performed the analysis. BA, FP and ST compiled the figures. BA and ST wrote the manuscript.

## ACKNOWLEDGEMENTS

We are grateful to Spyros Artavanis-Tsakonas, Chris Q. Doe, James Posakony, Yuh-Nung Jan, Jonas Muhr, the Developmental Studies Hybridoma Bank at the University of Iowa, and The Bloomington Drosophila Stock Center for sharing antibodies, fly lines, DNAs and advice. We thank Chris Q. Doe, Francois Schweisguth and Simon Sprecher for valuable comments on the manuscript. Annika Starkenberg, Carolin Jonsson and Helen Ekman provided excellent technical assistance. This work was supported by the Swedish Research Council, the Knut and Alice Wallenberg Foundation, the Swedish Cancer Foundation, and the University of Queensland, to ST.

---

**Supplemental Figure 1**

**Notch signalling regulates Wor and SoxN.**

(A-O) Expression of Dpn, Ase, Wor and *SoxN-GFP* in control, *neur^1^/neur^11^* and *AS-C* mutants, as well as *da-Gal4>UAS-l’sc* misexpression embryos, at St11. (A-C) Excessive numbers of NBs are generated in *neur^1^/neur^11^* mutants, whereas fewer NBs are produced in proneural deficiency embryos (AS-C), as evident by Dpn staining. (A-J) Ase and Wor expression shows similar expression profile, and co-localizes with Dpn in NBs, in both mutant and misexpression embryos (magnified panels). (K-O) In control, *SoxN-GFP* has a broader expression pattern than Dpn, Ase and Wor, being expressed both in NBs and in neighbouring ectodermal cells. *SoxN-GFP* expression responds in a complex manner to *neur^1^/neur^11^* and *Notch^55e11^* mutants (see Figure 2). *SoxN-GFP* is downregulated in *AS-C* and upregulated in *da-Gal4>UAS-l’sc* embryos. (P) Quantification of NB numbers inside the large dashed rectangles in (A-E) (Student’s two-tailed T-test; ***p≤0.001; **p≤0.01; n≥3 embryos; mean+/-SD). (Q-R) Quantification of Wor levels in NBs, and SoxN-GFP levels in lateral NBs, in stripes and in between stripes (Student’s two-tailed T-test; ***p≤0.001; n≥30 sections, n≥3 embryos; mean+/-SD). Scale bars: whole embryo = 50μm, magnified panels = 10μm.

**Supplemental Figure 2**

**Notch activates Crb expression.**

(A-F) Crb expression is decreased in Notch pathway mutants (*neur^1^/neur^11^* and *Notch^55e11^*) and increased in Notch pathway activated embryos (*da-Gal4>UAS-NICD* and *da-Gal4>UAS-m8^CK2^*). Boxes depicted on whole embryos are magnified below; dotted lines are shown as orthogonal views below. In control, NBs (Dpn+ cells) are observed delaminated from the overlying neurectodermal cell layer. In contrast, in both *neur* and *Notch* mutants, NBs are formed already in the overlying neurectodermal cell layer. (G-J) Quantification of Crb levels (Student’s two-tailed T-test; ***p≤0.001; n≥30 sections; n>3 embryos; mean+/-SD). Scale bars: whole embryo = 50μm, magnified panels = 10μm. Stage 11.

**Supplemental Figure 3**

**Proneural genes regulate Crb expression.**

(A-D) Crb expression is downregulated in both proneural mutants (AS-C) and misexpression (*da-Gal4>l’sc*). Small boxes and dotted lines on the whole embryo are visualized below as magnified panels and orthogonal views, respectively. (E-F) Quantification of Crb levels (fluorescence intensity; Student’s two-tailed T-test; ***p≤0.001; n≥30 sections, n≥3 embryos; mean+/-SD). Scale bars: whole embryo = 50μm, magnified panels = 10μm. Stage 11.

**Supplemental Figure 4**

***crb* regulates Wor and SoxN expression.**

(A-P) Expression of Dpn, Ase, Wor, SoxN-GFP and Crb in control, *crb* mutant (*crb^11A22^/crb^Df^*) and *crb* misexpression (*da-Gal4>UAS-crb*) embryos, at St11. Ase and Wor expression co-localizes with Dpn, in both mutant and misexpressing embryos (magnified panels below). Wor expression is reduced in *crb* mutants, but unaffected in *crb* misexpression. In control, *SoxN-GFP* has a broader expression pattern than Dpn, Ase and Wor, being expressed both in NBs and in neighbouring cells. *SoxN-GFP* expression is downregulated in both *crb^11A22^/crb^Df^* mutant and *da-Gal4>UAS-crb* misexpression embryos. (Q-R) Quantification of Wor levels in NBs, and SoxN-GFP levels in NBs, in stripes and in between stripes (fluorescence intensity; Student’s two-tailed T-test; ***p≤0.001; **p≤0.01; *p≤0.05; n≥30 sections, n≥3 embryos; mean+/-SD). Scale bars: whole embryo = 50μm, magnified panels = 10μm.

**Supplemental Table 1**

**Previously published DNA-binding data mining.**

DNA-binding analysis indicates extensive interplay between the Notch pathway, *SoxN, wor*, and EMT genes.

**Supplemental Table 2**

**RNA-seq analysis.**

RNA-seq analysis of co-misexpressed NB factors (*da>UAS-Quad* and *da>UAS-Quad^D^*) indicate extensive regulatory interplay between the Notch pathway, *SoxN* and wor, as well as the EMT genes.

