## Supplemental Data for "*Drosophila* Neuroblast Selection Gated by Notch, Snail, SoxB and EMT Gene Interplay"

### Supplemental Figure 1

#### Notch signalling regulates Wor and SoxN

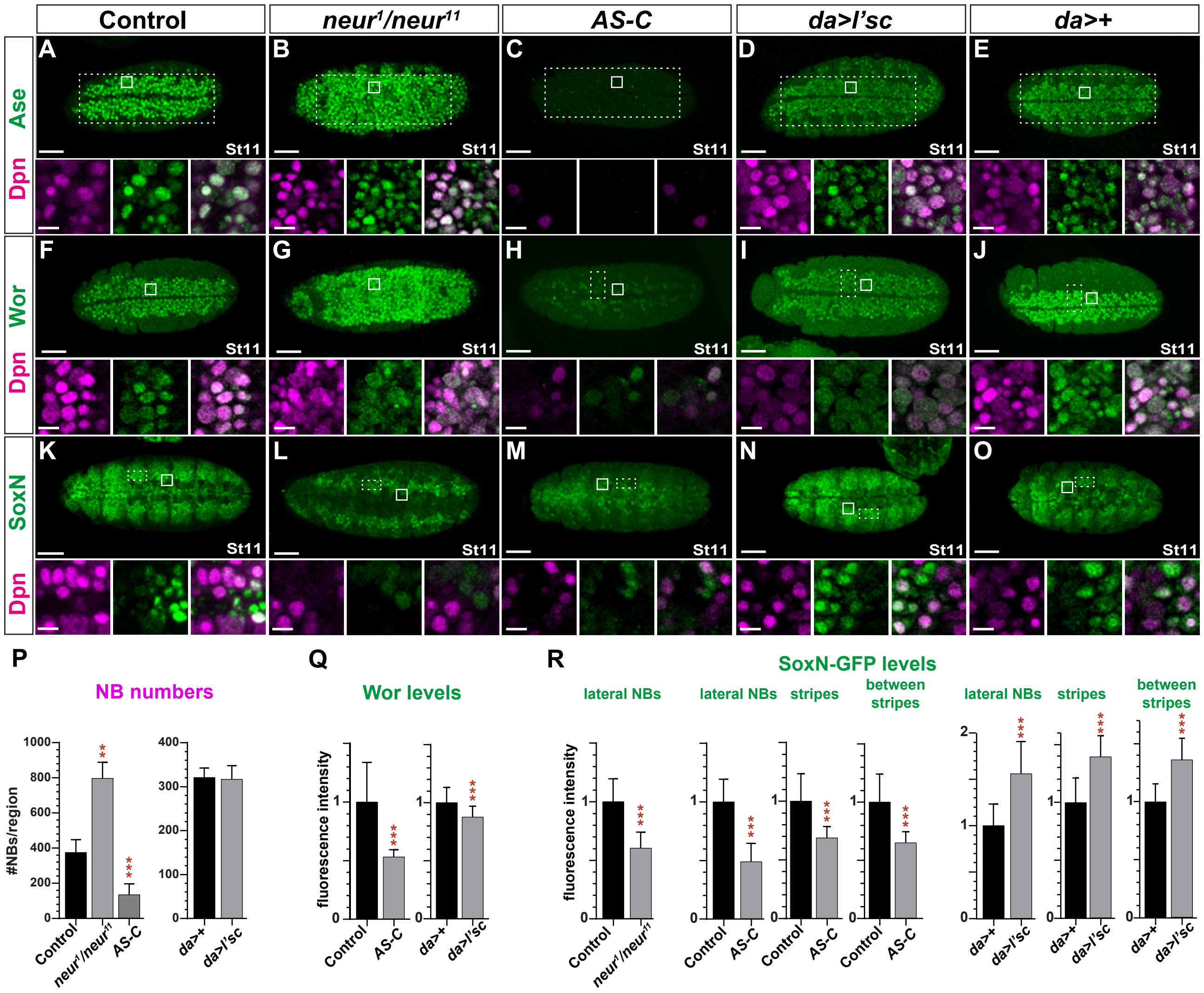

### Supplemental Figure 2

#### Notch activates Crb

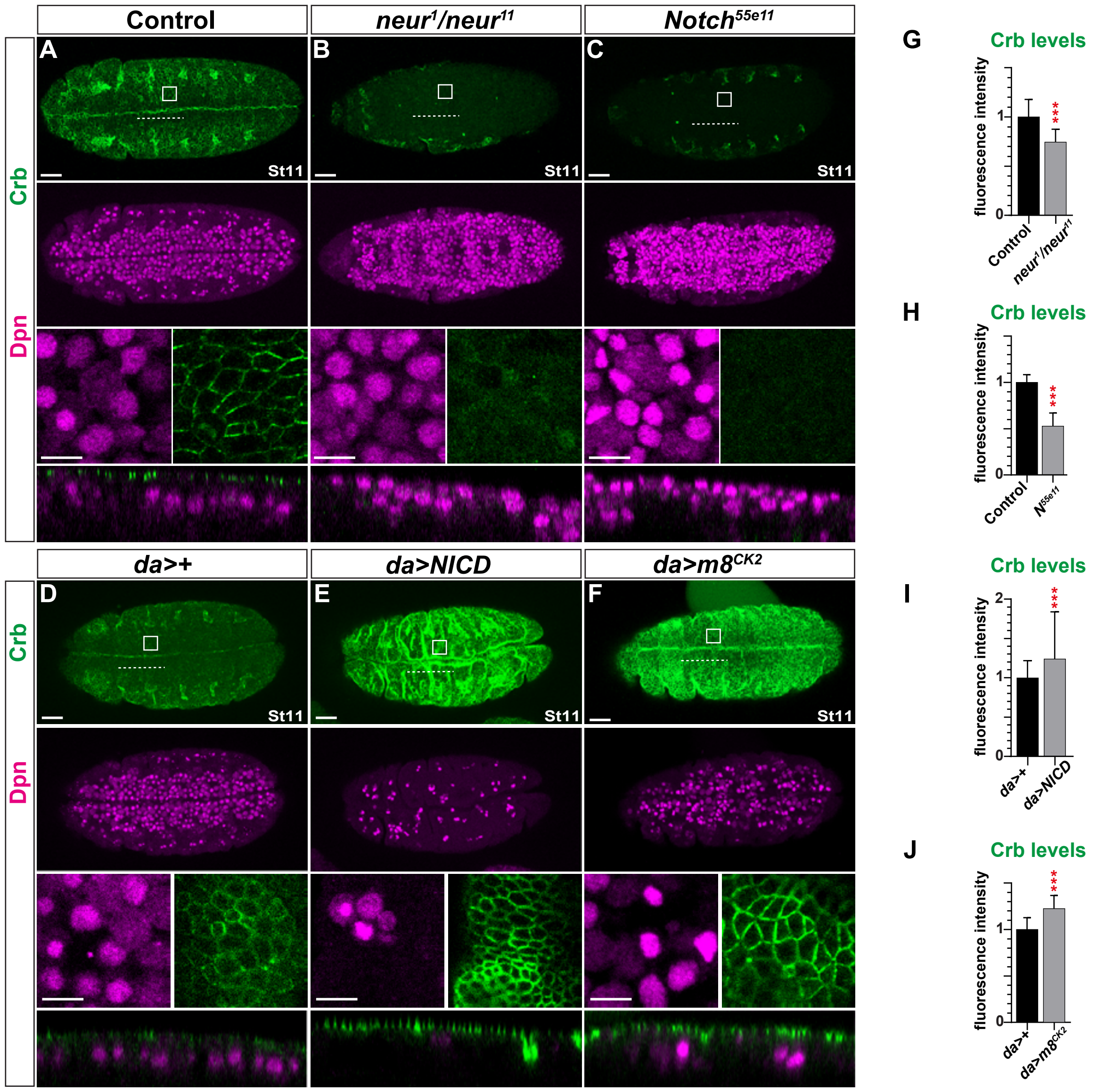

### Supplemental Figure 3

#### Proneural genes regulate Crb expression

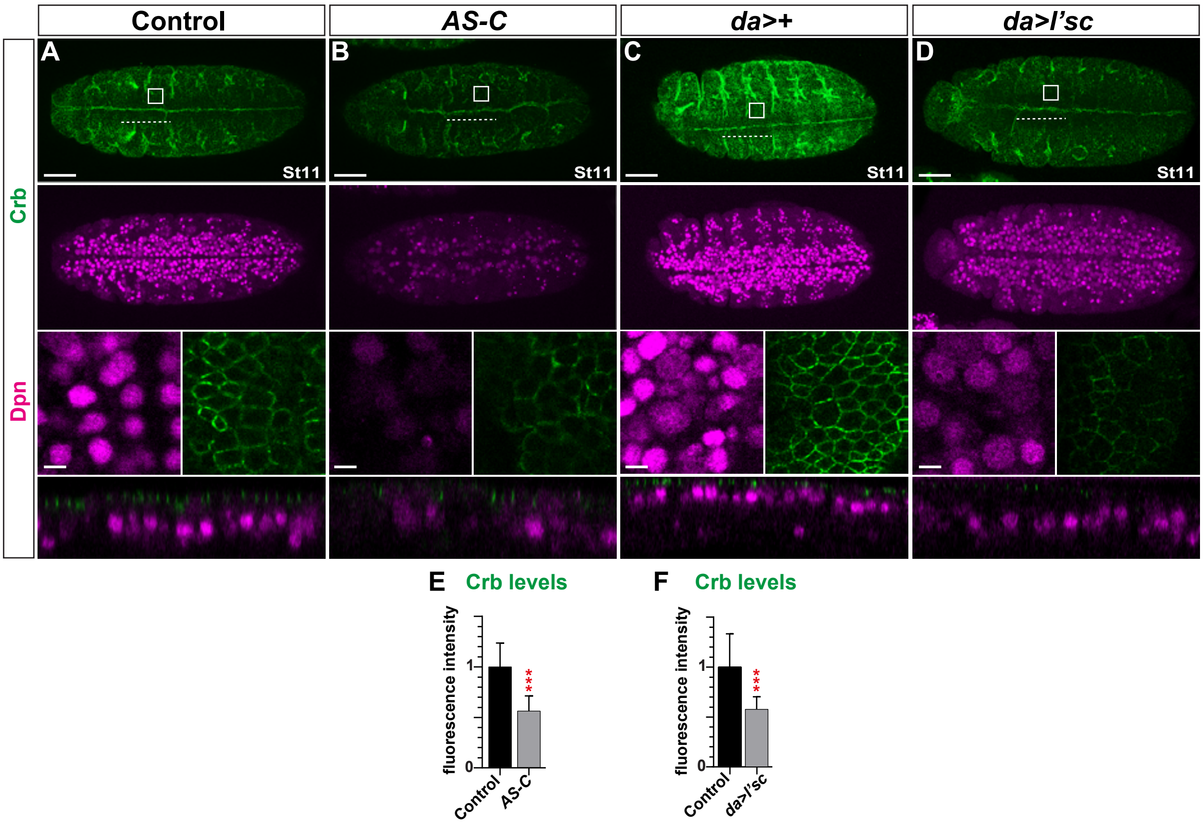

### Supplemental Figure 4

#### *crb* regulates Wor and SoxN expression

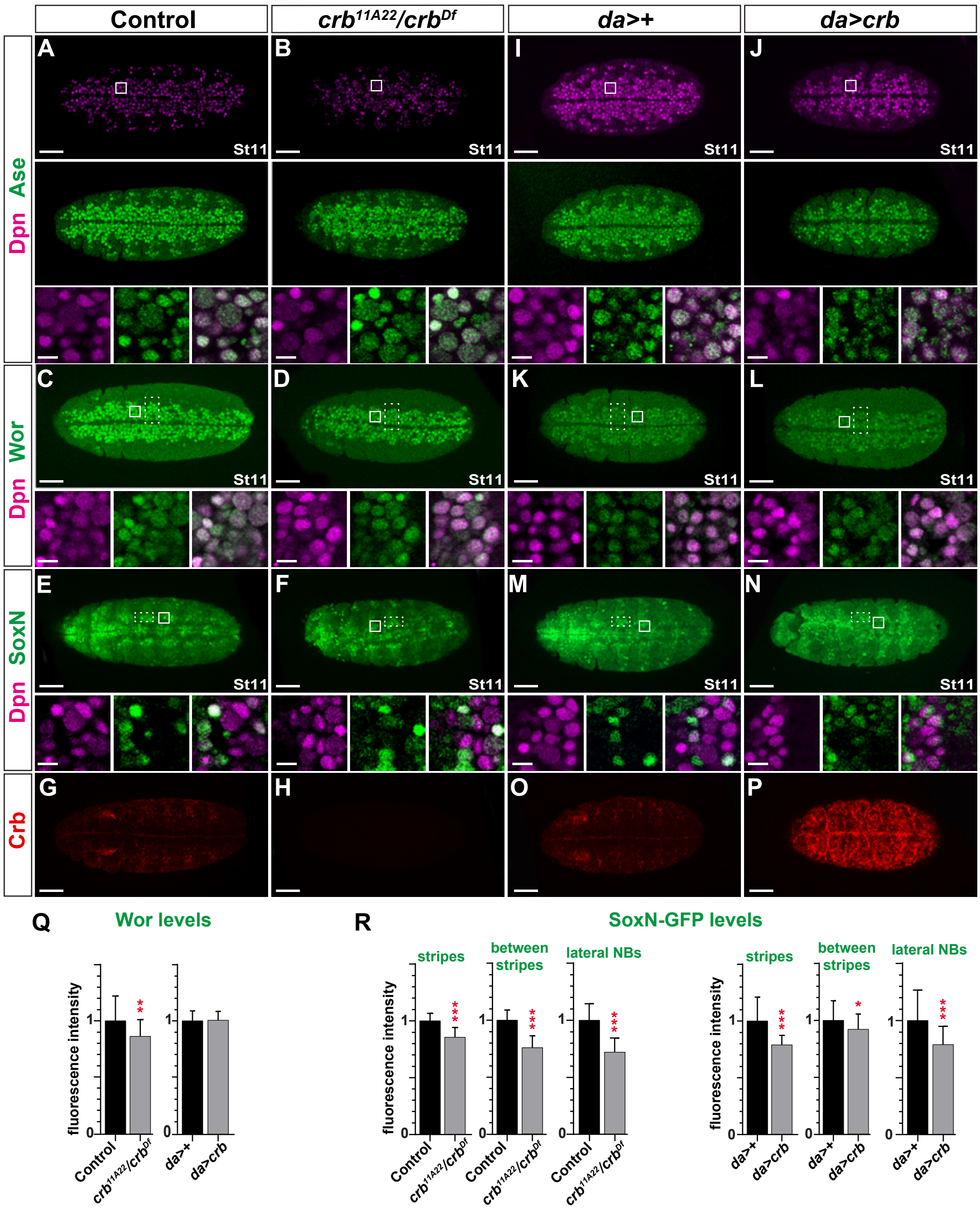

**Table S1, Related to Figure 1. Genes ≥2FC by whole embryo RNA-seq**

| | | Notch pathway | | | | | | | | Snail and SoxB genes | | | | | | EMT genes | | | | | Asymmetric NB genes | | | | | Par Complex | | | | Scribble Complex | | |
|---|---|---|---|---|---|---|---|---|---|---|---|---|---|---|---|---|---|---|---|---|---|---|---|---|---|---|---|---|---|---|---|---|
| Reference | Genotype | *E(spl)* | *Dl* | *neur* | *ac* | *sc* | *l'sc* | *ase* | | *wor* | *sna* | *esg* | *SoxN* | *D* | | *crb* | *sdt* | *Cad99C* | *Patj* | | *mira* | *insc* | *pros* | *pon* | | *baz (par3)* | *par-6* | *aPKC* | | *scrib* | *dlg1* | *l(2)gl (lgl)* |
| Bahrampour et al. 2017 | *da>Quad* | *m8* | | | | | | | | | | | | | | | | | | | | | | | | | | | | | | |
| | *da>Quad$^{D}$* | *m8* | | | | | | | | | | | | | | | | | | | | | | | | | | | | | | |

≥2FC Upregulated

>2FC Downregulated

Transgenic driven upregulation

<2FC

**Table S2, Related to Figure 1. Previously published DNA-binding.**

| | | | Notch pathway | | | | | | | | | Snail and SoxB genes | | | | | EMT genes | | | | Asymmetric NB genes | | | | Par Complex | | | Scribble Complex | | |
|---|---|---|---|---|---|---|---|---|---|---|---|---|---|---|---|---|---|---|---|---|---|---|---|---|---|---|---|---|---|---|
| **Genome-wide studies** | **Method** | **Protein** | *E(spl)* | *Dl* | *neur* | *Su(H)* | *N* | *ac* | *sc* | *l'sc* | *ase* | *wor* | *sna* | *esg* | *SoxN* | *D* | *crb* | *sdt* | *Cad99C* | *Patj* | *mira* | *insc* | *pros* | *pon* | *baz (par-3)* | *par-6* | *aPKC* | *scrib* | *dlg1* | *l(2)gl (lgl)* |
| ModEncode (www.modencode.org)(Negre et al., 2011) | ChIP-seq | D | ■ | ■ | ■ | ■ | ■ | | ■ | ■ | ■ | ■ | ■ | ■ | | ■ | ■ | ■ | ■ | ■ | | ■ | ■ | | ■ | ■ | ■ | | ■ | |
| Embryo (on gene body, or within 10kb of gene) | ChIP-seq | Sna | ■ | ■ | ■ | ■ | ■ | ■ | ■ | ■ | ■ | | ■ | ■ | ■ | ■ | ■ | ■ | ■ | ■ | ■ | | ■ | ■ | ■ | ■ | ■ | ■ | ■ | ■ |
| Southall and Brand, 2009, embryo | Dam-ID | Ase | ■ | ■ | ■ | ■ | ■ | | ■ | ■ | ■ | ■ | ■ | | | ■ | | | | | ■ | | ■ | | ■ | | ■ | | | |
| | Dam-ID | Sna | ■ | ■ | ■ | ■ | ■ | ■ | ■ | ■ | | | ■ | ■ | ■ | ■ | | ■ | | | | | ■ | | ■ | | | ■ | ■ | |
| Aleksic et al., 2013, embryo | DamID | D | ■ | ■ | ■ | | ■ | | ■ | | | ■ | ■ | ■ | ■ | ■ | | ■ | ■ | | ■ | ■ | ■ | | ■ | ■ | ■ | ■ | ■ | ■ |
| Bivik et al., 2016, embryo | ChIP-seq | Su(H) | ■ | ■ | ■ | ■ | | ■ | ■ | ■ | ■ | ■ | ■ | ■ | ■ | ■ | ■ | ■ | ■ | ■ | ■ | ■ | ■ | | ■ | ■ | ■ | ■ | ■ | ■ |
| On gene body, or within 10kb of gene | DamID | E(spl)m5 | ■ | ■ | ■ | ■ | ■ | ■ | ■ | ■ | ■ | ■ | ■ | ■ | ■ | ■ | ■ | ■ | ■ | ■ | ■ | ■ | ■ | | ■ | ■ | ■ | ■ | ■ | ■ |
| | DamID | E(spl)m8 | ■ | ■ | ■ | ■ | ■ | ■ | ■ | ■ | | ■ | ■ | ■ | ■ | ■ | ■ | ■ | ■ | ■ | ■ | ■ | ■ | | ■ | ■ | ■ | ■ | ■ | ■ |
| Krejci et al., 2009, DmD8 and Kc cells | ChIP-CHIP | Su(H) | ■ | | ■ | | ■ | | | | | | | | | | | | | | | | | | | | | | | |
| Bernard et al., 2010, DmD8 cells | ChIP-CHIP | Su(H) | ■ | | ■ | | ■ | | | | | | | | | | | | | | | | | | | | | | | |
| Djiane et al., 2013, wing discs | ChIP-CHIP | Su(H) | ■ | ■ | ■ | | ■ | | | | | | | | | | | | | | | | | | | | | | | |
| Housden et al., 2013, DmD8 cells | ChIP-CHIP | Su(H) | ■ | | | | ■ | | | | | | | | | | | | | | | | | | | | | | | |
| | | | ■ | | | | ■ | | | | | | | | | | | | | | | | | | | | | | | |
| **Gene-focused studies** | | | | | | | | | | | | | | | | | | | | | | | | | | | | | | |
| Krejci et al., 2007, S2 and DmD8 cells | ChIP-PCR | Su(H) | ■ | | | | | | | | | | | | | | | | | | | | | | | | | | | |
| Lecourtois and Schweisguth, 1995 | EMSA | Su(H) | ■ | | | | | | | | | | | | | | | | | | | | | | | | | | | |
| Nellesen et al., 1999 | EMSA | Su(H) | ■ | | | | | | | | | | | | | | | | | | | | | | | | | | | |
| Cave et al., 2005 | EMSA | Su(H) | ■ | | | | | | | | | | | | | | | | | | | | | | | | | | | |
| Bailey and Posakony, 1995 | EMSA | Su(H) | ■ | | | | | | | | | | | | | | | | | | | | | | | | | | | |
| Liu and Posakony, 2014 | EMSA | Su(H) | | | | ■ | | | | | | | | | | | | | | | | | | | | | | | | |
| Barolo et al., 2000 | EMSA | Su(H) | | | | ■ | | | | | | | | | | | | | | | | | | | | | | | | |
| Miller and Posakony, 2018 | EMSA | Sc | | | ■ | | | | | | | | | | | | | | | | | | | | | | | | | |
| Bailey and Posakony, 1995 | EMSA | Ac | ■ | | | | | | | | | | | | | | | | | | | | | | | | | | | |
| Oellers et al, 1994 | EMSA, footprint | L'Sc | ■ | | | | | ■ | | | | | | | | | | | | | | | | | | | | | | |
| Tietze et al, 1992 | EMSA | E(spl)m8 | ■ | | | | | | | | | | | | | | | | | | | | | | | | | | | |
| Oellers et al, 1994 | EMSA, footprint | E(spl)m5, m8 | ■ | | | | | ■ | | | | | | | | | | | | | | | | | | | | | | |
| Van Douren et al, 1994 | EMSA | E(spl)m7 | | | | | | ■ | | | | | | | | | | | | | | | | | | | | | | |
| **Gene location** | | | **3R** | **3R** | **3R** | **2L** | **X** | **X** | **X** | **X** | **X** | **2L** | **2L** | **2L** | **2L** | **3L** | **3R** | **X** | **3R** | **3L** | **3R** | **2R** | **3R** | **X** | **X** | **X** | **2R** | **3R** | **X** | **2L** |
